# Unlocking the Distinct Roles of the Three Mammalian VDAC Isoforms in Mitochondrial Respiration and Cancer Cell Metabolism

**DOI:** 10.1101/2025.02.20.639106

**Authors:** Megha Rajendran, William M. Rosencrans, Wendy Fitzgerald, Diana Huynh, Bethel G. Beyene, Baiyi Quan, Julie Hwang, Nina A. Bautista, Tsui-Fen Chou, Sergey M. Bezrukov, Tatiana K. Rostovtseva

**Affiliations:** Eunice Kennedy Shriver National Institute of Child Health and Human Development, NIH, Bethesda, Maryland, USA 20892; Division of Biology and Biological Engineering, California Institute of Technology, Pasadena, California, USA 91125

## Abstract

The Voltage Dependent Anion Channel (VDAC) is the most ubiquitous protein in the mitochondrial outer membrane. This channel facilitates the flux of water-soluble metabolites and ions like calcium across the mitochondrial outer membrane. Beyond this canonical role, VDAC has been implicated, through interactions with protein partners, in several cellular processes such as apoptosis, calcium signaling, and lipid metabolism. There are three VDAC isoforms in mammalian cells, VDAC 1, 2, and 3, with varying tissue-specific expression profiles. From a biophysical standpoint, all three isoforms can conduct metabolites and ions with similar efficiency. However, isoform knockouts (KOs) in mice lead to distinct phenotypes, which may be due to differences in VDAC isoform interactions with partner proteins. To understand the functional role of each VDAC isoform within a single cell type, we created functional KOs of each isoform in HeLa cells and performed a comparative study of their metabolic activity and proteomics. We found that each isoform KO alters the proteome differently, with VDAC3 KO dramatically downregulating key members of the electron transport chain (ETC) while shifting the mitochondria into a glutamine-dependent state. Importantly, this unexpected dependence of mitochondrial function on the VDAC3 isoform is not compensated by the more ubiquitously expressed VDAC1 and VDAC2 isoforms. In contrast, VDAC2 KO did not affect respiration but upregulated ETC components and decreased key enzymes in the glutamine metabolic pathway. VDAC1 KO specifically reduced glycolytic activity linked to decreased hexokinase localization to mitochondria. These results reveal non-redundant roles of VDAC isoforms in cancer cell metabolic adaptability.

## Introduction

Voltage-dependent anion channel (VDAC), an outer mitochondrial membrane (MOM) protein, is crucial for the exchange of small ions such as calcium and water-soluble metabolites such as ATP, ADP, and NADH between cytosol and mitochondria. VDAC, a monomeric weakly anion-selective ß-barrel channel, not only conducts but also effectively regulates their fluxes across MOM. In vitro studies with VDAC reconstituted into the planar lipid membranes showed that one of the mechanisms of such regulation is VDAC’s characteristic and evolutionarily conserved voltage-dependent gating when under applied voltage, the channel moves from high-conducting unique “open state” passable for ATP, to low-conducting so-called “closed states” that are impermeable to ATP^1^ but more permeable for calcium^2^. Recent studies in vitro have revealed that metabolite and calcium fluxes through VDAC can also be regulated by VDAC complexation with various cytosolic proteins such as dimeric tubulin and α-synuclein^3-5^. Furthermore, VDAC is known to be involved in the regulation of calcium signaling through the formation of complexes with calcium transporters on the endoplasmic reticulum (ER)^6^, sarcoplasmic reticulum (SR)^7,8^, and lysosomes^9^. In addition, VDAC is implicated in apoptosis through its interaction with several pro-and anti-apoptotic proteins^10-16^. This highlights the intricate role of VDAC in regulating metabolism, calcium signaling, and cell death, which is further complicated by the isoform-specific differences in VDAC interactome and function.

There are three VDAC isoforms in mammals: VDAC1, 2, and 3. VDAC1 and 2 diverged from VDAC3, which is considered the oldest isoform^17,18^. All three isoforms show high sequence similarity (∼ 70%) and form similar anion-selective voltage-gating channels in vitro based on single-channel electrophysiology experiments^19,20^. However, mice knockout (KO) studies suggest distinct roles for each VDAC isoform, with VDAC2 KO resulting in development delays and embryonic lethality^21,22^. VDAC1 KO mice exhibit mild bioenergetics defects^23^ and VDAC3 KO results in male infertility^24^. The latter was attributed to each isoform’s tissue-specific role and expression level since VDAC3 is the highest expressed isoform in testis. However, VDAC3 KO also results in mitochondrial dysfunction in heart muscles, where it is the least expressed isoform. VDAC1 and 3 demonstrate different affinities for interaction with two identified VDAC cytosolic regulators - tubulin and α-synuclein^20^. Dimeric α/ß-tubulin is the building block of microtubules, and α-synuclein is a neuronal protein implicated in Parkinson’s disease (PD). These two abundant cytosolic proteins have no structural, functional, or genetic similarity but possess a common feature—disordered and negatively charged C-terminal tails that act as effective VDAC blocking domains by being reversibly captured by the VDAC pore and reversing its selectivity to cationic by a similar mechanism^25^. VDAC1 binds tubulin and α-synuclein with a 10-100 times higher affinity than VDAC3^20^. In addition, reconstituted VDAC isoforms differ in calcium selectivity, with VDAC3 having a higher calcium preference than VDAC1^26^. De Stefani et al. showed similar results in cells with VDAC3 overexpression, resulting in increased mitochondrial calcium uptake, while VDAC1 may play a role in apoptotic calcium signaling^27^. VDAC2 has recently emerged as an important regulator of mitochondrial calcium uptake, especially in the cardiomyocytes^28,29^. Interestingly, although both VDAC1 and VDAC2 are implicated in apoptosis, each of them interacts with a different set of Bcl-2 family proteins: VDAC1 preferably interacts with anti-apoptotic Bcl2, Bcl-xL^14,30^, and VDAC2 with pro-apoptotic Bak and Bax proteins^21,22,31^.

VDAC1 and 2 are the most abundant isoforms in most human tissues, and VDAC1 is the only one whose crystal structure has been solved^32-34^. Therefore, rather naturally, VDAC1 isoform is the most studied in vitro and in vivo, followed by VDAC2. VDAC3 isoform, in contrast, is the least expressed in vivo (except for testis) and the least biophysically and biochemically studied in vitro isoform. Despite the initial publication by Colombini’s and Craigen’s labs in 1999^19^, where VDAC3’s ability to form typical voltage-gated ion channels has been demonstrated, the following publications often reported that VDAC3 does not form voltage-gated channels^35^ or even that it cannot form channels at all^36^. Only recently, the record was straightened out in Queralt-Martin et al. work^20^, where it was unambiguously demonstrated that reconstituted VDAC3, similar to VDAC1 and 2, forms typical anion-selective voltage-gated channels. One of the differences between isoforms is the number of cysteines, with VDAC2 and VDAC3 having seven and nine cysteines, respectively, and VDAC1 having only two cysteines^37,38^. Such a drastic difference focused the studies of VDAC3 on its role in oxidative stress^39^, where it was shown that VDAC3 KO in HAP1 cells promoted the accumulation of free radicals dependent on the cysteines, thus suggesting a role for VDAC3 cysteines in countering ROS-induced oxidative stress.

Early studies using mice knockout models demonstrated VDAC isoform-specific differences in mitochondrial functions. VDAC1 KO resulted in a universal decrease in the activity of electron transport chain (ETC) complexes, but VDAC3 KO specifically affected mitochondrial function in the heart and sperm^40^. VDAC1 KO in MEF cells showed decreased respiration and glycolytic capacity^41^. However, until recently, VDAC was mostly ignored in mitochondrial bioenergetics studies as it was historically considered a “molecular sieve” in MOM, except for the studies focused on VDAC1’s role in MOM permeabilization at the earlier stages of apoptosis^42,43^ and its tentative involvement in the mitochondrial Permeability Transition Pore (PTP)^44,45^. Recently, Magri et al. showed that VDAC1 KO diminished complex I-linked respiration in HAP1 cells due to decreased respiratory reserves^46^. However, Yang et al. reported no change in respiration but showed increased glycolysis due to VDAC1 KO in H9c2 cells^47^. Beyond the mentioned discrepancies, a comprehensive systematic study comparing the role of each VDAC isoform in mitochondrial respiration and metabolism in the same cell type is lacking.

To understand the mechanism of differences in isoform function in metabolism, we systematically investigated the role of each VDAC isoform knockout using CRISPR-Cas9 mediated gene editing in HeLa cells. We found that VDAC1 KO affected glycolysis, while VDAC2 KO did not measurably affect glycolysis or mitochondrial respiration. Surprisingly, it was VDAC3 KO that decreased mitochondrial respiration. Proteomic studies revealed increased expression of mitochondrial proteins upon VDAC2 KO in contrast to extensive downregulation of mitochondrial proteins in VDAC3 KO cells. Overexpression of glutaminases in VDAC3 KO cells, necessary for mitochondrial glutamine metabolism, results in a dependence of these cells on glutamine. We demonstrate that VDAC3, the least expressed isoform, is crucial for mitochondrial function to fuel high-energy demand. Overall, the data reveals non-redundant roles of VDAC isoforms in cancer cell metabolic adaptability.

## Materials and methods

### Cell culture

HeLa cells (CCL-2) were purchased from ATCC (American Type Culture Collection). HeLa cells with CRISPR-Cas9 knockout cell lines of VDAC1, VDAC2, and VDAC3 were generated by Synthego. The cells were grown in DMEM (Gibco, 15607) supplemented with 10% fetal bovine serum (Gibco, 10437) at 37 °C and 5% CO_2_.

### Cell proliferation assay

Cells were seeded at 2,500 cells per well in 96-well plates (ThermoFisher Scientific, 165305), and cell count was measured every 24 hours. Cells were washed with PBS and fixed with 4% PFA (Electron Microscopy Sciences, 15710) containing 25 µM Hoechst 33342 (ThermoFisher Scientific, 62249) for 15 mins. Cells were washed with PBS and imaged using a DAPI filter in BioTek Lionheart FX automated microscope.

CellTiter 96® AQ_ueous_ One Solution Cell Proliferation Assay (MTS) (Promega, G3582) was performed based on manufacturer protocol. Cells were seeded at 2,500 cells in a 96-well plate and incubated at 37 °C in a 5% CO_2_ incubator for 72 hours. 20 µL CellTiter 96® AQ_ueous_ One Solution Reagent was added to 100 µL media per well and incubated for 1 hour at 37 C in a 5% CO_2_ incubator. The absorbance was measured at 490 nm using a CLARIOstar plate reader.

### Seahorse metabolic flux assay

Cells were seeded at 10,000-15,000 cells/well in Seahorse XFp PDL miniplates (Agilent, 103721) or XFe96/XF Pro PDL cell culture microplates (Agilent, 103798). They were incubated overnight at 37 °C and 5% CO_2_. The following day, the media was replaced with Seahorse XF DMEM assay media (Agilent, 103575) supplemented with glucose, pyruvate, and glutamine based on the manufacturer’s protocol for cell mitochondrial stress (Agilent, 103010), glycolysis stress (Agilent, 103020), and mitochondria fuel flex (Agilent, 103260) assays. The assays were performed on Seahorse XF HS mini or XF Pro analyzers (Agilent) according to the manufacturer’s protocol. Hoechst 33342 (ThermoFisher Scientific, 62249) was added to the last port to stain nuclei for cell count and imaged using Lionheart FX or Cytation5 (BioTek).

### Colocalization of hexokinase to mitochondria

Cells were seeded at 25,000 cells/well in µ-Slide 8 well chambered coverslip (Ibidi, 80807) and grown at 37 °C and 5% CO_2_ overnight. Cells were transfected with HK1-GFP (Addgene, 21917) or HK2-GFP (Addgene, 21920) along with Omp25-mCherry (Addgene, 157758) using Lipofectamine™ Stem transfection reagent (ThermoFisher Scientific, STEM00008) according to the manufacturer’s protocol. The cells were washed and imaged in Fluorobrite DMEM (Gibco, A18967) with a 20x/0.75 N.A. objective in Leica TCS SP8 microscope. GFP was excited with a 488 nm laser, and emission was captured using a 500–582 nm filter, and mCherry was excited with a 587 nm laser and emission was captured using a 592-784 nm filter. Images were optimized for contrast and brightness, and Pearson’s correlation coefficient was measured using the BIOP JACoP plugin in ImageJ (NIH).

### Western blot

After harvesting, cell pellets were resuspended in 150 µL lysis buffer (50 mM Tris-HCl pH 8.0, 150 mM NaCl, 1% Triton X-100 with protease inhibitor tablet (Pierce), and incubated on ice for 10 min with occasional vortex. Samples were centrifuged at 15,000 rpm at 4 °C for 10 min, and 120 µL of the supernatant was transferred into a new 1.5 mL tube. Total soluble protein concentrations were measured using the Bradford reagent (Bio-Rad, 5000006). After that, 40 µL of 4x Laemmli sample buffer (Bio-Rad, 161–0774) containing 0.1 M DTT (Cytiva, 17-1318-02) was mixed with the samples and heated for 5 min at 95 °C. Equal amounts of protein samples were loaded and separated using 4–20% Mini-PROTEAN TGX precast gels (Bio-Rad, 456– 1096) and transferred to nitrocellulose membranes using Trans-Blot Turbo Transfer System (Bio-Rad, 170–4155). Membranes were blocked with 5% w/v nonfat milk prepared in TBST buffer, incubated with primary antibodies overnight at 4C, washed with 5% milk (three times for 10 min each), incubated with proper secondary antibodies for two h at room temperature, and washed with TBST buffer (three times for 5 min each). The blots were imaged using an ECL reagent (MilliporeSigma, WBKLS0500) and the ChemiDoc MP Imaging System (Bio-Rad). Blot densities were analyzed using Image Lab 6.0.1 software (Bio-Rad).

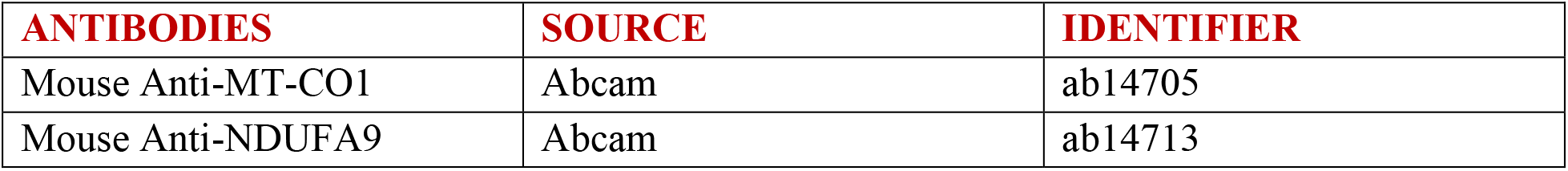
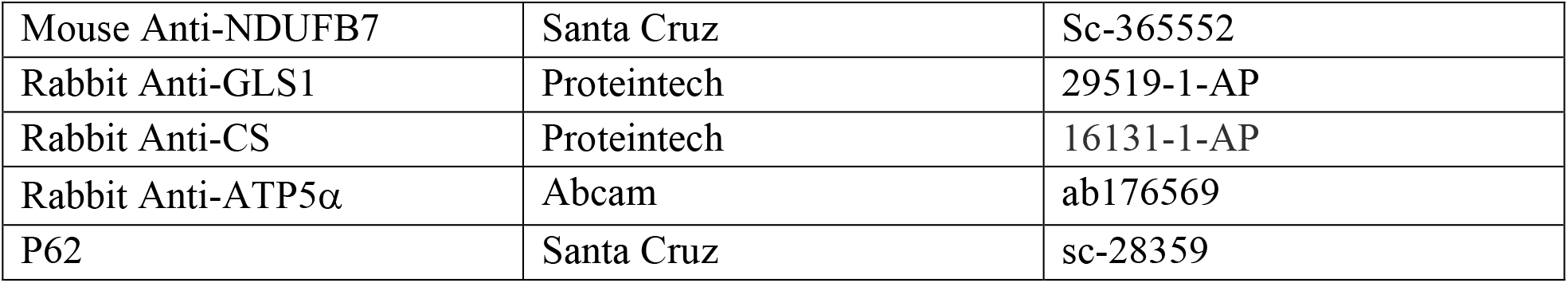

| ANTIBODIES | SOURCE | IDENTIFIER |
| --- | --- | --- |
| Mouse Anti-MT-CO1 | Abcam | ab14705 |
| Mouse Anti-NDUFA9 | Abcam | ab14713 |
| Mouse Anti-NDUFB7 | Santa Cruz | Sc-365552 |
| Rabbit Anti-GLS1 | Proteintech | 29519-1-AP |
| Rabbit Anti-CS | Proteintech | 16131-1-AP |
| Rabbit Anti-ATP5α | Abcam | ab176569 |
| P62 | Santa Cruz | sc-28359 |

### Label-free proteomics

1.2 million cells were harvested with trypsin, quenched with PBS, spun down to remove the trypsin, and resuspended and spun down in ice-cold PBS. The supernatant was removed, and LC-MS samples were prepared using the Thermo EasyPep Mini MS Sample Prep Kit (ThermoFisher Scientific, A4006) according to the manufacturer’s instructions. Samples were then resuspended in 0.1% formic acid (ThermoFisher Scientific, 85178) solution, and peptide concentration was tested using the Pierce Quantitative Fluorometric Peptide Assay (ThermoFisher Scientific, 23290). LC-MS/MS experiments were performed by loading a 500 μg sample onto an EASY-nLC 1000 (ThermoFisher Scientific) connected to an Orbitrap Eclipse Tribrid mass spectrometer (ThermoFisher Scientific). Peptides were separated on an Aurora UHPLC Column (25 cm x 75 μm, 1.6 μm C18, AUR2-25075C18A, Ion Opticks) with a flow rate of 0.4 mL/min and for a total duration of 131 min. The gradient was composed of 3% Solvent B for 1 min, 3–19% B for 72 min, 19–29% B for 28 min, 29–41% B for 20 min, 41–95% B for 3 min, and 95–98% B for 7 min. Solvent A consists of 97.8% H2O, 2% acetornitrile, and 0.2% formic acid, and solvent B of 19.8% H2O, 80% ACN, and 0.2% formic acid. MS1 scans were acquired with a range of 400–1600 m/z in the Orbitrap at 120 k resolution. The maximum injection time was 50 ms, and the AGC target was 2 3 105. MS2 scans were acquired using quadrupole isolation mode and higher-energy collisional dissociation (HCD) activation type in the Iontrap. The isolation window was 1.4 m/z, collision energy was 35%, maximum injection time was 35 ms, and the AGC target was 1 3 104. Other global settings were set to the following: ion source type, NSI; spray voltage, 2500 V; ion transfer tube temperature, 300 °C. Method modification and data collection were performed using Xcalibur software (ThermoFisher Scientific).

### Quantification and statistical analysis

Proteomic analysis was performed using Proteome Discoverer 2.4 (PD 2.4, ThermoFisher Scientific) software, the Uniprot human database, and SequestHT with Percolator validation. Percolator FDRs were set at 0.001 (strict) and 0.01 (relaxed). Peptide FDRs was set at 0.001 (strict) and 0.01 (relaxed), with medium confidence and a minimum peptide length of 6. Carbamidomethyl (C) was set as a static modification; oxidation (M) was set as a dynamic modification; acetyl (protein N-term), Met-loss (Protein N-term M) and Met-loss + acetyl (Protein N-term M) were set as dynamic N-Terminal modifications. Protein abundance normalization was performed relative to the total peptide amount. Differential Expression analysis was performed with media using a custom Python module following the user guide 7 or one-tailed Student’s t-test with PD 2.4. Principal Component Analysis (PCA) analyses were conducted with custom Python code. Volcano plots were generated in R using the Tidy Proteomics and Enhanced Volcano plot packages. Venn diagram was plotted using Python3. Gene ontology analysis was performed using g:Profiler (website (https://biit.cs.ut.ee/gprofiler/gost). Other statistical analyses were carried out by one-tailed Student’s t-test or one-way ANOVA using Prism 7.0. Three independent biological replicates were used. p values < 0.05 are reported as statistically significant and are depicted as follows throughout the manuscript: * p < 0.05, ** p < 0.01, **** p < 0.0001.

## Results

### Characterization of HeLa VDAC isoform knockout cells

Functional Knockouts (KOs) of each VDAC isoform were generated using CRISPR-Cas9 in HeLa cells. The indels were confirmed by sequencing (Supplementary Figure S1), and the KO was confirmed by western blot (Supplementary Figure S2A). HeLa cells were chosen as they express characteristic levels of VDAC isoforms found in most human cell types. Although different proteins may respond differently to mass spectrometry (MS) quantitation, considering the sequence similarity amongst the VDAC isoforms, we postulated that the relative abundances for VDAC isoforms are proportional to their copy numbers. Through MS observation, VDAC1 is the most expressed isoform (∼50%), followed by VDAC2 (∼30%), and VDAC3 (∼18%) is the least expressed isoform in HeLa cells (Supplementary Figure S2B). The metabolic activity for all VDAC KO clones was investigated by measuring NADH (Supplementary Figure S3A), FAD (Supplementary Figure S3B) fluorescence using flow cytometry, and ATP production rates (Supplementary Figure S4) using Seahorse XF Pro Analyzer. While VDAC2 KO clones do not affect NADH and FAD levels, VDAC1 and VDAC3 KO showed some variability. VDAC1 KO consistently decreased glycolytic ATP production, but mitochondrial ATP production rate was variable (Supplementary Figure S4A). VDAC3 KO results in different glycolytic and mitochondrial ATP production rates (Supplementary Figure S4C). Such variability between clones is intriguing and suggests a compensatory mechanism. We chose to focus on the representative clones with altered mitochondrial function VDAC1 (G8), VDAC2 (C9), and VDAC3 (E5) to understand the isoform-specific role of this mitochondrial outer membrane (MOM) protein in metabolism in HeLa cells.

Conventional thinking would assume that the overexpression of other VDAC isoforms might compensate for the loss of one isoform. Using the MS results, the expression level of each VDAC isoform in each of the KOs could be directly compared. We found only mild changes in VDAC isoform expression in response to specific KO. VDAC1 KO cells show a ∼20% decrease in VDAC2 expression, and VDAC2 KO increases VDAC3 expression (∼40%), but these changes do not compensate for the KO of each isoform (Figure 1A). The KO cells do not show appreciable difference in cell morphology (Supplementary Figure S5A), but VDAC2 and VDAC3 KOs decrease the growth rate of HeLa cells by 30-40% in 72 hours (Supplementary Figure S5 B and C). Cell proliferation was also measured using the CellTiter 96® AQ_ueous_ One Solution Cell Proliferation Assay. The assay contains an MTS tetrazolium compound that is reduced by NADPH or NADH in metabolically active cells into a colored formazan product with absorbance at 490 nm. While this assay is widely used to measure the number of viable cells, the absorbance depends on the metabolic activity of cells. Interestingly, only VDAC3 KO significantly decreases the absorbance (∼40-55%) (Figure 1B), which suggests that VDAC3 KO specifically affects the metabolic activity in HeLa cells.

**Figure 1.**
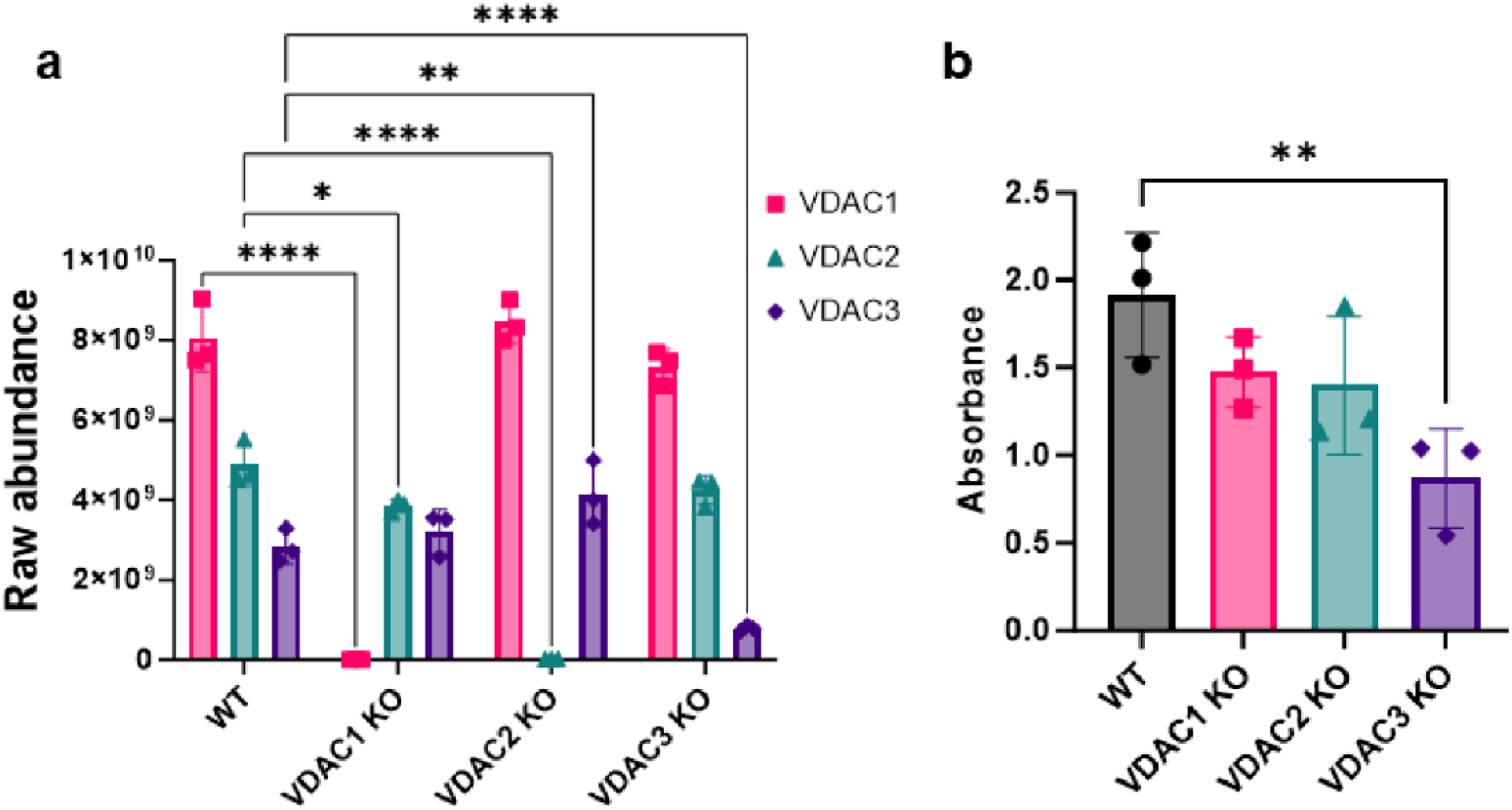
VDAC3 KO decreases metabolic activity in HeLa cells. a) The bar graph represents the raw abundance of VDAC1 (pink), VDAC2 (teal), and VDAC3 (purple) in WT, VDAC1 KO, VDAC2 KO, and VDAC3 KO HeLa cell lines measured using mass spectrometry (MS). The ∼30% expression of VDAC3 in VDAC3 KO despite western blot confirmation complete KO (Supplementary Figure S2B) may correspond to the expression of a shorter transcript since the two VDAC3 peptides detected by MS correspond to the C-terminal of VDAC3 (Supplementary MS table). b) The bar graph shows the metabolic activity of WT, VDAC1 KO, VDAC2 KO, and VDAC3 KO cells 72 hours post seeding measured using CellTiter 96® AQueous one solution cell proliferation assay. The symbols represent data from 3 independent experiments, and error bars indicate the standard deviation from the mean. Significance was tested using one-way ANOVA followed by the Dunnett post hoc test (*p< 0.05, **p<0.01, ****p<0.0001)

To decipher the role of each VDAC isoform in metabolism, we compared the metabolic activity using a Seahorse XF analyzer.

### VDAC1 KO affects basal glycolysis in HeLa cells

To examine the role of each isoform in glycolysis, we performed a glycolysis stress test, which measures the changes in Extracellular Acidification Rate (ECAR) due to increased glycolytic activity in cells. Figure 2A shows the ECAR trace in WT and KO cells upon the addition of glucose, oligomycin, and 2-deoxyglucose (2-DG). Basal glycolysis is calculated by subtracting non-glycolytic acidification after adding 2-DG (red box) from the increase in ECAR upon adding glucose (blue box). Only VDAC1 KO decreases basal glycolysis. Surprisingly, VDAC3 KO resulted in the complete loss of glycolytic capacity in HeLa cells, even though basal glycolysis was unaffected (Figure 2C). Glycolytic capacity (Figure 2A green box) measures the ability of cells to increase glycolysis upon the inhibition of OXPHOS with oligomycin, which may be linked to mitochondrial dysfunction in VDAC3 KO cells.

**Figure 2.**
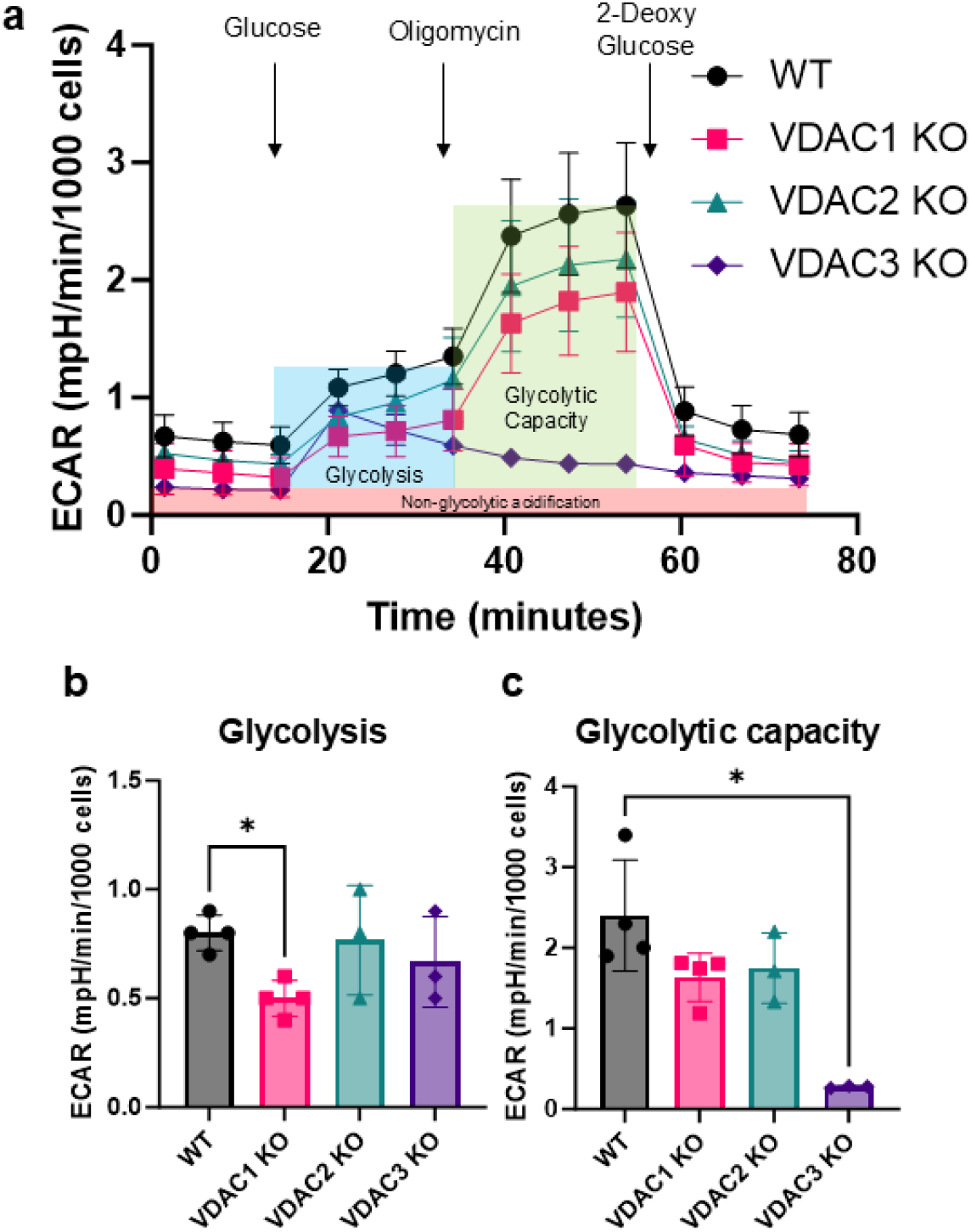
VDAC1 KO decreases glycolysis in HeLa cells. a) Traces of extracellular acidification rate (ECAR) upon the addition of glucose, oligomycin, and 2-deoxyglucose for WT (black circle), VDAC1 KO (pink square), VDAC2 KO (teal triangle), and VDAC3 KO (purple diamond) cells. Glycolysis (blue box) and glycolytic capacity (green box) are calculated after subtraction from non-glycolytic acidification (red box). Bar graphs comparing glycolysis (b) and the glycolytic capacity (c) between WT (gray), VDAC1 KO (pink), VDAC2 KO (teal), and VDAC3 KO (purple) HeLa cells. Data from 3-4 independent experiments are represented in the bar graph, and the error bars indicate the standard deviation from the mean. The symbols represent data from independent experiments. Significance was tested using one-way ANOVA followed by the Dunnett post hoc test (*p < 0.05).

### VDAC3 KO decreases mitochondrial respiration in HeLa cells

The mitochondrial stress test was performed to determine the effect of VDAC isoform KOs on mitochondrial respiration. The Oxygen Consumption Rate (OCR) was measured under basal conditions and after adding oligomycin, FCCP, and rotenone/antimycin-A (Figure 3A). VDAC1 KO and VDAC2 KO showed no significant change in respiration compared to WT HeLa WT cells (Figure 3B). However, VDAC3 KO significantly decreased basal and maximal respiration (Figure 3B). Interestingly, VDAC3 KO results in a complete loss of spare respiratory capacity (SRC) (Figure 3C), which suggests that HeLa cells with VDAC3 KO are unable to meet increased energy demand during acute cellular stress. SRC depends on the availability of mitochondrial substrates, suggesting a role for VDAC3, the least expressed isoform, in the uptake of mitochondrial substrates in HeLa cells.

**Figure 3.**
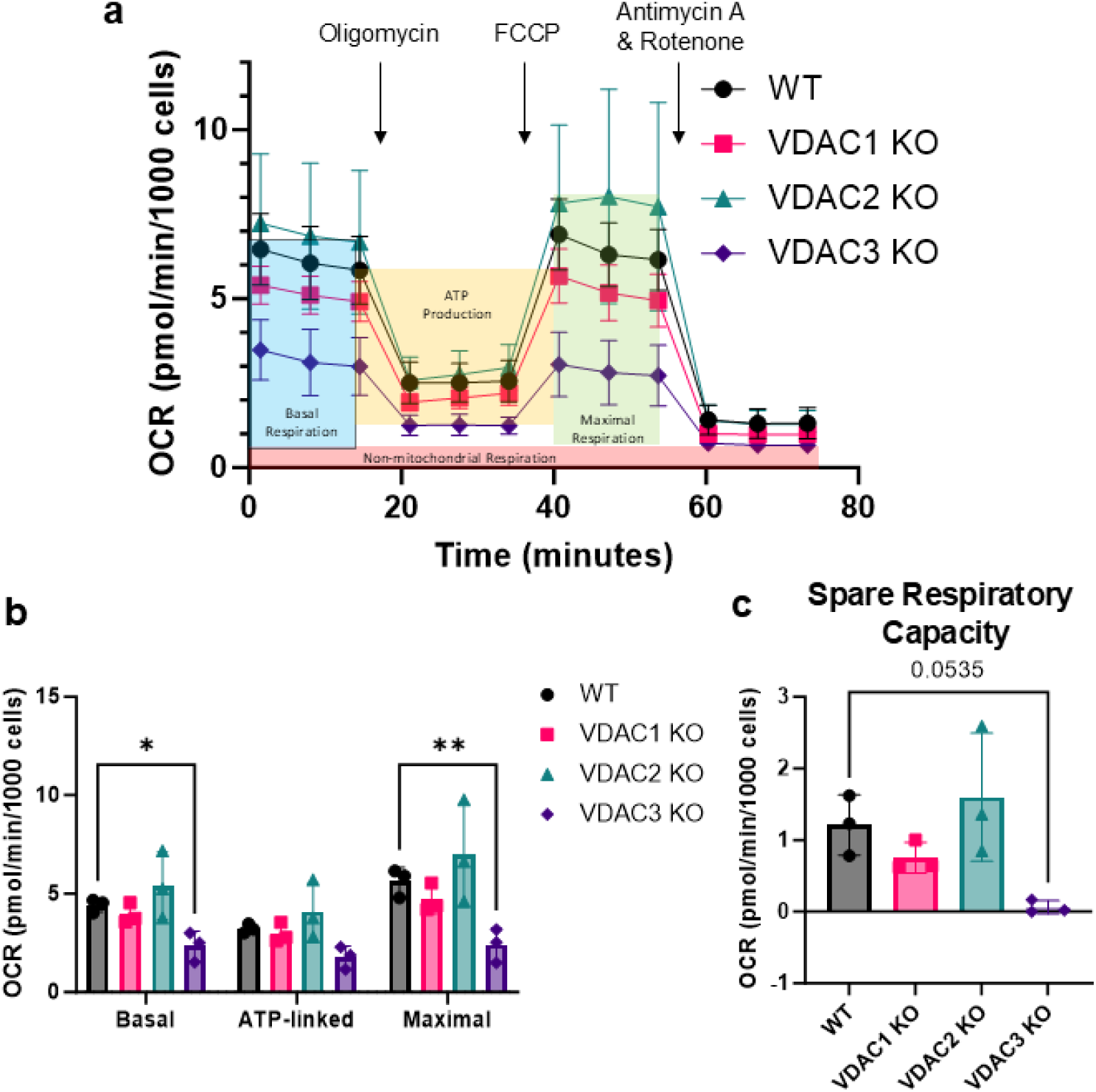
VDAC3 KO decreases mitochondrial respiration in HeLa cells, resulting in the complete loss of spare respiratory capacity. a) Trace of oxygen consumption rate (OCR) upon the addition of oligomycin, FCCP, and Antimycin A/Rotenone for WT (black circle), VDAC1 KO (pink square), VDAC2 KO (teal triangle), and VDAC3 KO (purple diamond) cells. Non-mitochondrial respiration (red box) is subtracted from basal (blue box), ATP-linked (yellow box), and maximal (green box) respiration for analysis. b) Bar graphs comparing basal, ATP-linked, maximal respiration, and SRC (c) in WT (gray), VDAC1 KO (pink), VDAC2 KO (teal), and VDAC3 KO (purple) HeLa cells. Data from three independent experiments (symbols) are averaged, and the error bar indicates the standard deviation from the mean. Significance was tested using one-way ANOVA followed by the Dunnett post hoc test (*p < 0.05, **p < 0.01).

### VDAC shows isoform-specific changes in cellular proteome

To understand the molecular basis of these striking phenotypes of VDAC isoforms KOs at the level of protein expression, label-free proteomics was performed on each of the KO cell lines. Using liquid chromatography coupled with tandem MS, we identified and quantified over 9,500 unique proteins across all four cell lines: WT, VDAC1 KO, VDAC2 KO, and VDAC3 KO.

Most proteins (9,168) were identified in all studied cell lines, enabling direct comparison without the need for excessive imputation (Figure 4A). Expression data were subjected to a median normalization ^48^. The principal component analysis (PCA) demonstrated tight clustering of each of the sample replicates within each cell type compared to different genetic backgrounds (Figure 4B). Each cell line is separated into distinct regions of the principal component plot of the PC1/PC2 plot, with the WT cells remaining directly between each VDAC KO. These data indicate that each VDAC isoform KO generally leads to a unique change in the cellular proteome.

**Figure 4.**
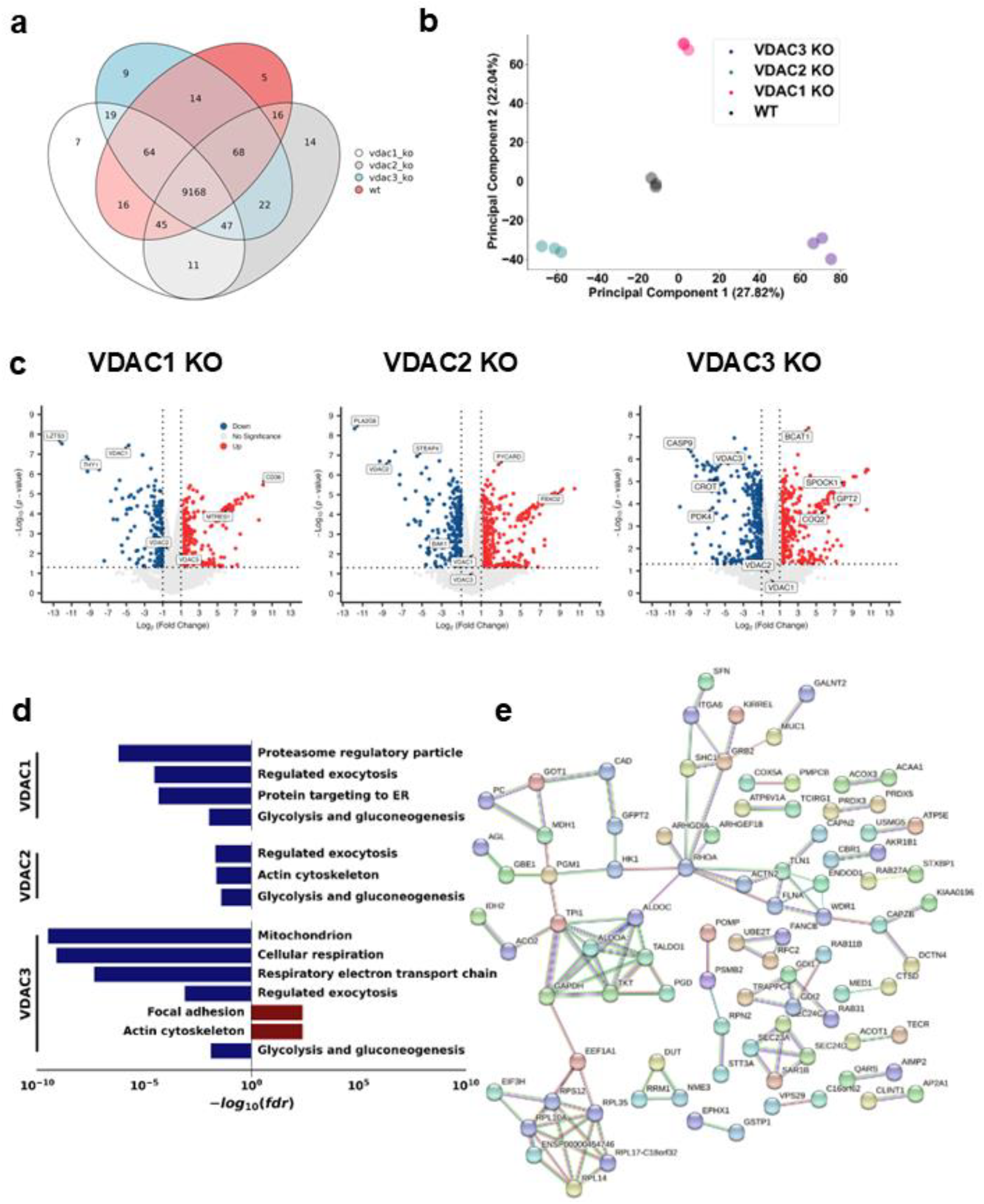
Total proteomics reveals isoform-specific proteome remodeling in HeLa cells. a) Venn diagram detailing the number of unique proteins identified in each sample group. b) PCA plot demonstrating the stratification of each VDAC isoform KO sample compared to WT cells along PC1 and PC2. c) Volcano plots of each VDAC isoform KO compared to WT cells. Proteins with > 1-fold absolute change and p-value < 0.05 are colored blue for downregulation and red for upregulation compared to the WT. Proteins of interest are labeled. d) Representative GO Term analysis showing pathways significantly downregulated upon each VDAC isoform (blue) or upregulated (red) compared to the WT. e) STRING analysis of glycolysis and gluconeogenesis proteins downregulated in all VDAC isoform KOs compared to WT.

To determine the proteins that were most altered in each KO compared to the WT, differential protein expression analysis was performed. Focusing on those proteins with the most significant and largest magnitude of change between the different KOs, we observed alteration in several proteins hitherto not connected to VDAC in the existing literature. In the VDAC1 KO cells, the highest magnitude loss was of the transcription factor LZTS3 (Figure 4C). Among the most expressed proteins upon VDAC1 KO was the fatty-acid transporter CD36. In the VDAC2 KO cells, the highest magnitude loss was in the protein PLA2G6, a phospholipase, a gene for which loss of function results in a series of infantile neurodevelopmental disorders and Parkinson’s disease. PLA2G6 mutations are known to cause mitochondrial deficits, putatively from alterations in lipid homeostasis^49^. We also observed the loss of the metallo-reductase STEAP4, which is associated with mitochondrial dysfunction^50^. VDAC2 KO led to overexpression of the apoptosis and inflammasome-related protein PYCARD and the upregulation of the E3 ubiquitin ligase FBOX2 protein. Notably, VDAC2 KO led to a downregulation of Bak. VDAC2 is known to be essential for the import of the pro-apoptotic Bak into the MOM^21^. The downregulation of Bak may indicate a response to the reduced mitochondrial import of Bak in the absence of VDAC2.

Naghdi et al also showed the importance of VDAC2 in mitochondrial recruitment of Bak in HepG2 cells and hepatocytes ^51^. Interestingly, autophagy receptor p62 was downregulated in HeLa cells with VDAC3 KO. The western blot further confirmed that VDAC3 KO decreases p62 expression (Supplementary Figure S6). This result suggests either an alteration to autophagy pathways or oxidative stress pathways via KEAP1-Nrf2 due to the loss of VDAC3^52^. VDAC3 KO resulted in the loss of cell death and mitochondrial-related protein CASP9, as well as the metabolic protein PDK4. In VDAC3 KO, we observed upregulation of the spermatogenesis protein SPOCK1, consistent with VDAC3’s unique role in this developmental process ^24^. These data indicate that the majority of pathways altered by each VDAC KO are unique.

To investigate the pathways uniquely perturbed by each isoform, we performed unbiased gene ontology analysis of VDAC KO proteomics data to identify groups of pathways both conserved and altered uniquely by different isoform KOs (Figure 4D). Among pathways downregulated in all VDAC isoform KOs, the loss of glycolysis and gluconeogenesis-related proteins was observed. Proteins associated with these terms include hexokinase 1 (HK1) (Figure 4E). Two HK isoforms, HK1 and HK2, are suggested to regulate glycolysis through their interaction with mitochondria. We imaged the colocalization of HK1-GFP (Figure 5A) and HK2-GFP (Supplementary Figure S7A) with mitochondria labeled with Omp25-mCherry in HeLa WT and KO cells. Though all VDAC isoform KOs downregulate the expression level of HK1, only VDAC1 KO significantly decreases the localization of HK1(Figure 5B) and HK2 (Supplementary Figure S7B) to the mitochondria. Other lowered proteins in the glycolytic pathway included aldolases A and C, transaldolase, and GAPDH. We also observed a downregulation in proteins related to exocytosis in all KO cells (Figure 4D). Of pathways altered uniquely in each isoform, VDAC1 KO showed a reduction in proteasome regulation and ER protein trafficking. VDAC2 KO resulted in a minor downregulation of proteins related to the actin cytoskeleton, which is upregulated in the VDAC3 KO cells compared to WT, along with focal adhesion (Figure 4D). VDAC3 KO cells show the strongest effects on mitochondrially annotated proteins with significant losses of proteins involved in the ETC and oxidative respiration. Following these links to the modulation of mitochondrial pathways, we focused on the alterations to the mitoproteome.

**Figure 5.**
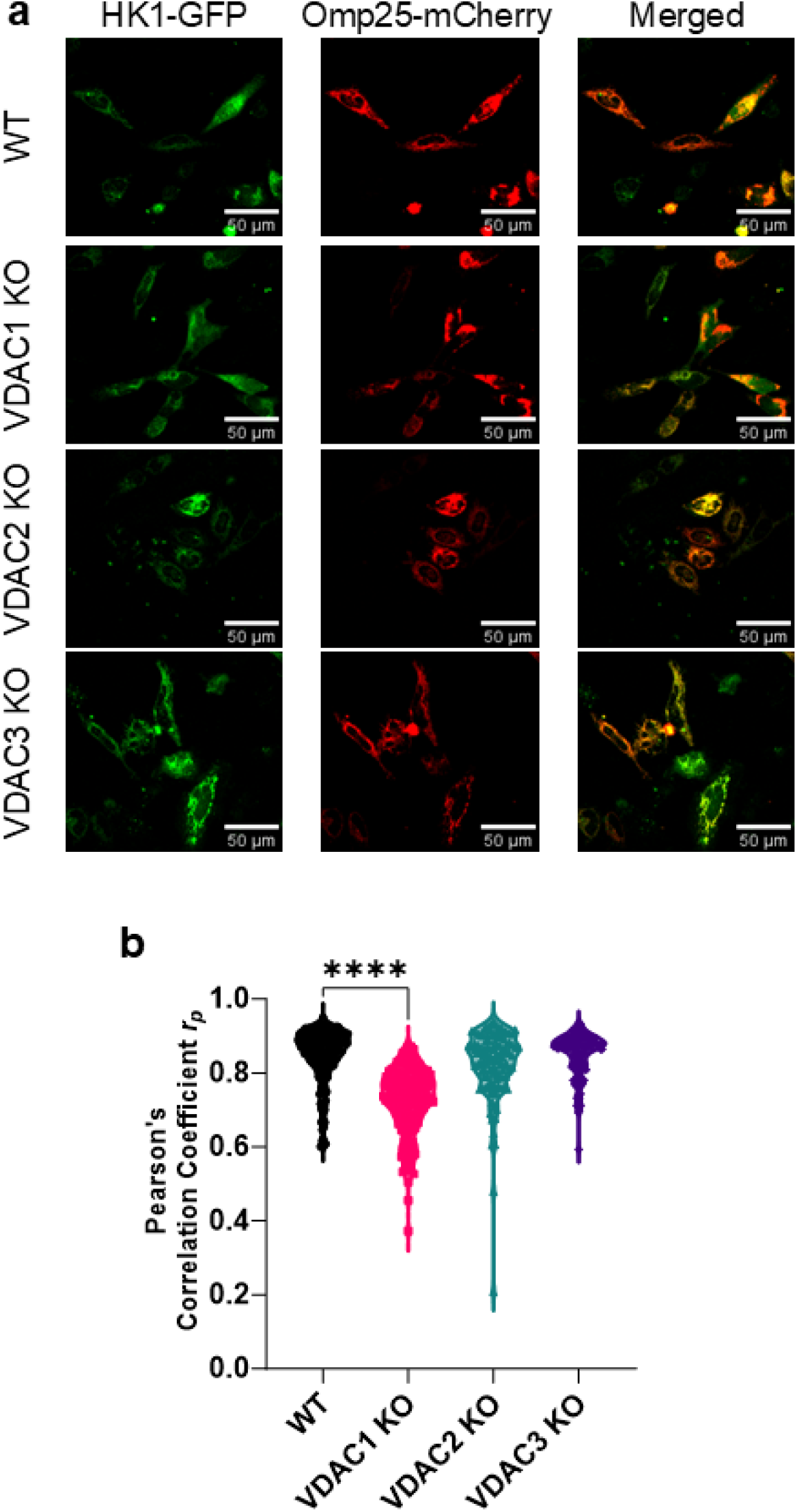
Mitochondrial localization of HK1 is dependent on VDAC1 isoform. a) Representative images of HK1-GFP (green) colocalization with Omp25-mCherry (red) shown in the merged image (yellow) for WT, VDAC1 KO, VDAC2 KO, and VDAC3 KO HeLa cells. b) Colocalization analysis of HK1 with Omp25 measured by Pearson’s correlation coefficient (PCC) shows a significant decrease in HK1 mitochondrial localization in VDAC1 KO. The symbols represent PCC for each cell. Significance was tested using one-way ANOVA followed by the Dunnett post hoc test (****p<0.0001).

### VDAC isoform KOs rewire the mitoproteome

To focus on the mitoproteome, we segmented those proteins annotated as mitochondrial in the Mitocarta3 database. All VDAC isoform KOs showed some alteration to mitochondrial proteins, with 26 proteins found downregulated in all VDAC KOs (Figure 6A). VDAC3 KO cells have 116 unique downregulated mitochondrial proteins, consistent with the GO term analysis. In contrast, VDAC3 KO cells had the least number of unique mitochondrial proteins upregulated at 32, compared to VDAC1 with 84 and VDAC2 with 96. Only 14 proteins were found upregulated across all VDAC KOs. These data support the notion that VDAC3 KO shows severe loss of mitochondrial proteins, whereas VDAC1 and VDAC2 KO appear to show milder alterations in mitochondrial proteins. The STRING analysis of mitochondrial proteins uniquely altered in each VDAC KO shows that VDAC3 KO cells have a large, downregulated cluster of proteins involved in the ETC (Figure 6B, red box), as described in the GO Term analysis. In contrast, VDAC2 KO cells show an upregulation of a cluster of ETC-related proteins. These results show that VDAC1 KO causes the least defined alteration to the mitochondrial proteome in HeLa cells despite VDAC1 being one of the most ubiquitous mitochondrial proteins.

**Figure 6.**
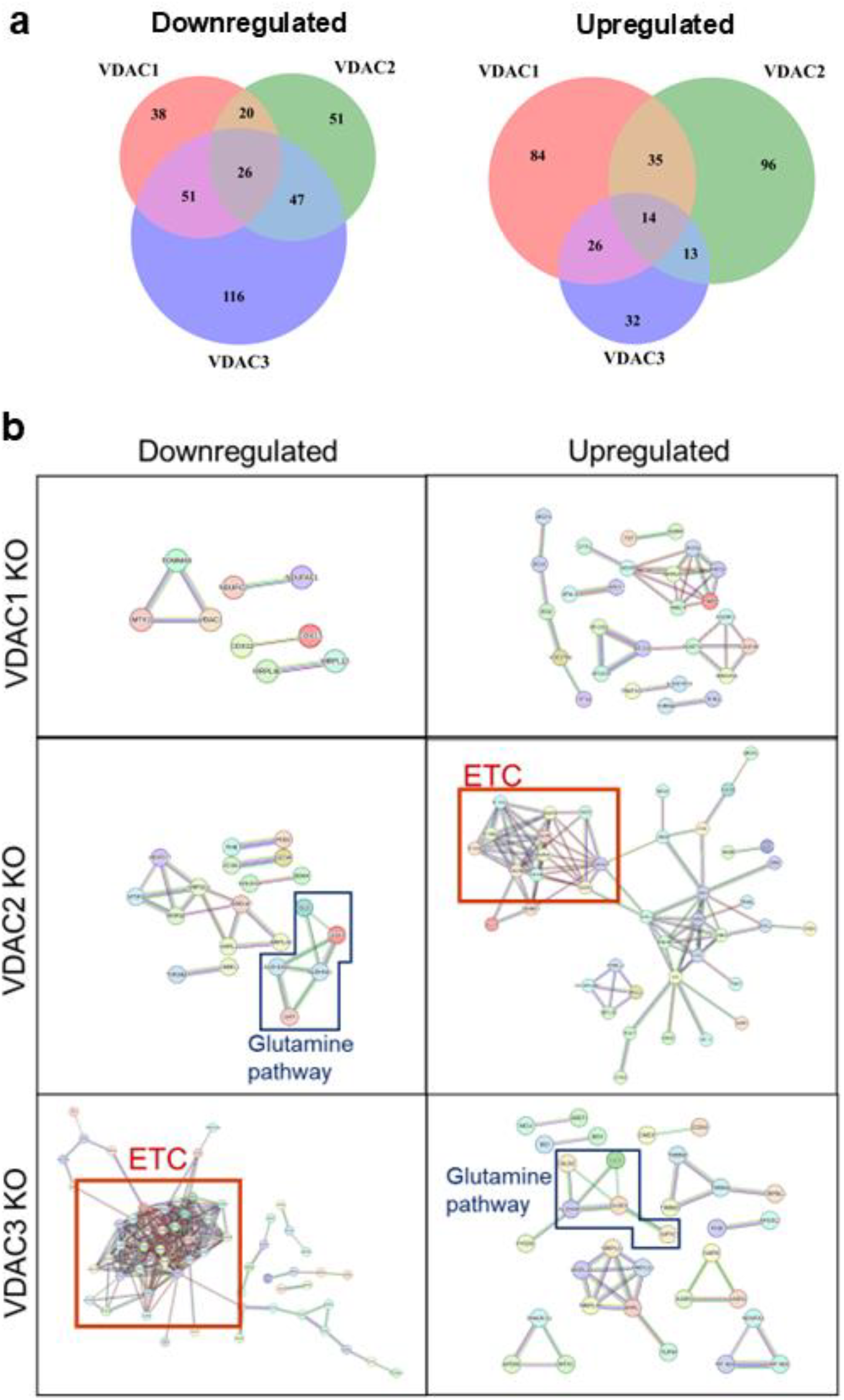
VDAC isoform KO leads to distinct mitochondrial proteome remodeling. a) Venn diagram showing mitochondrially annotated proteins found to be downregulated or upregulated in each of the three VDAC isoform KOs compared to WT. b) STRING analysis of mitochondrial pathways uniquely downregulated or upregulated in each VDAC isoform KO cell lines. Proteins involved in the electron transport chain (ETC) (red box) and glutamine pathway (blue box) are highlighted.

### VDAC3 KO leads to defects in the ETC

Figure 7A shows that VDAC3 KO causes a substantial loss in expression across most ETC components, whereas VDAC2 KO shows mostly upregulation in the mass-spectrometry results. In contrast, VDAC1 KO cells demonstrate the least perturbation in the ETC proteome. The proteomic results are further validated by a select number of these hits by western blot (Figure 7B, C). The proteins in complex I (NDUFA9), complex IV (MT-CO1), and complex V (ATP5A) are found to be consistently downregulated in the VDAC3 KO cells. In contrast, VDAC2 KO showed significant upregulation of ATP5A but downregulation of NDUFA9 and MT-CO1. In addition, the VDAC3 KO causes a slight, but not significant, defect in the expression of TCA enzyme citrate synthase (CS), whereas VDAC2 KO upregulated CS. The western blots show similar expression trends. VDAC3 KO causes the most severe defects, consistent with decreased mitochondrial respiration (Figure 3).

**Figure 7.**
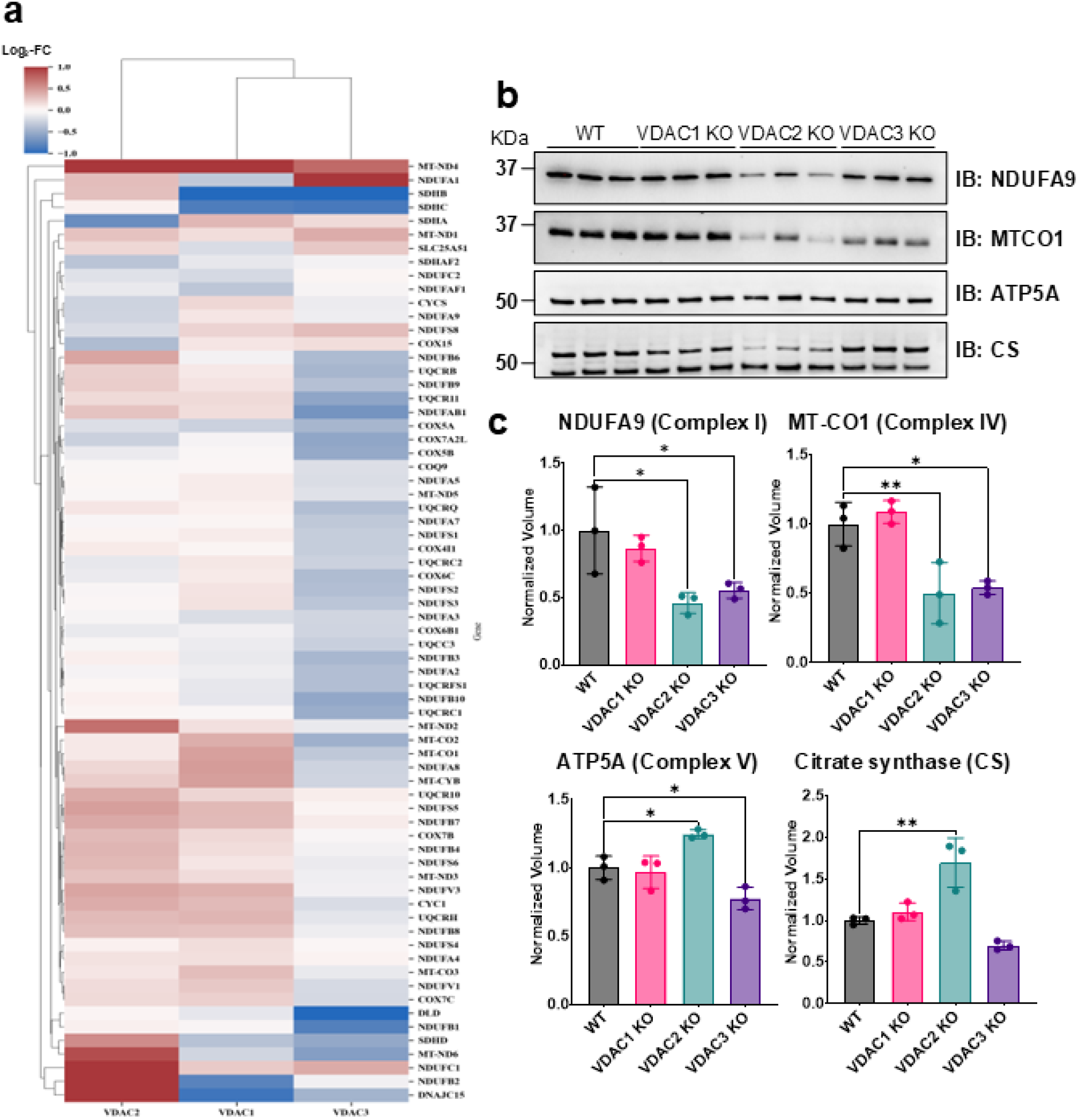
VDAC2 KO and VDAC3 KO result in widespread upregulation and downregulation of ETC components. a) Heatmap of altered proteins associated with the mitochondrial ETC. b) Western blot of selected ETC components across VDAC isoform KO cell lines and (c) the corresponding densitometry quantification. Data for three repeats are averaged in the bar graphs, and error bars indicate the standard deviation from the mean. The symbols represent data from each repeat. Significance was tested using one-way ANOVA followed by the Dunnett post hoc test (* p < 0.05, ** p < 0.01).

### VDAC3 KO leads to an increased dependence on glutamine metabolism

The STRING analysis of the mitochondrial proteome (Figure 6B) across VDAC KO lines shows an upregulation of glutamine pathway proteins in VDAC3 KO cells and a downregulation in VDAC2 KO (Figure 6B, blue box). To get insight into these surprising results, we investigated the expression of other proteins in the glutamine metabolic pathway across the VDAC KO cells.

We found that VDAC3 KO upregulates glutaminases (GLS1/2) and GLUD1/GPT2 (Figure 8A). GLS1 is a key mitochondrial enzyme that converts glutamine into glutamate, which GLUD1/GPT2 then converts into α-ketoglutarate to feed the TCA cycle. Western blotting further confirms the upregulation of GLS1 in VDAC3 KO cells (Figure 8B). These data indicate that VDAC3 KO cells rewire the metabolic program pathways, increasing reliance on glutamine for mitochondrial respiration. Notably, the upregulation of glutaminase is a hallmark of certain cancers, such as ovarian, breast, and colorectal cancer, resulting in their dependence on glutamine ^53^. Targeting glutaminase in these cancers has been a popular target of therapeutic intervention ^54^. To determine whether these alterations have functional effects on the metabolic state of the VDAC KO cells, we performed a mitochondrial fuel flex test using a Seahorse XF analyzer to determine their dependencies on glucose, glutamine, and fatty acid to fuel mitochondrial respiration. In this assay, a series of metabolic inhibitors is used to separate the contribution of each fuel source to respiration. UK5099 blocks the mitochondrial pyruvate carrier (MPC) to inhibit the glucose oxidation pathway; etomoxir inhibits carnitine palmitoyltransferase 1A (CPT1A), which transports long chain fatty acid into mitochondria inhibiting long chain fatty acid oxidation pathway; and BPTES inhibits GLS1, which converts glutamine to glutamate inhibiting glutamine oxidation pathway (Figure 8C). The relative fuel dependence can be estimated by measuring the oxygen consumption rate in the presence or absence of the fuel pathway inhibitors. WT HeLa cells have equal dependence on glucose and glutamine with a small contribution from fatty acid oxidation (Figure 8D). We found that VDAC3 KO has a significantly increased (∼30%) dependence on glutamine metabolism, demonstrating that the overexpression of GLS and other glutamine pathway proteins leads to a functional change in metabolic phenotype (Figure 7D). There is also an increased dependence on fatty acid metabolism in the VDAC3 KO cells. VDAC1 KO cells show a 37% decrease in glucose dependence compared to WT, while VDAC2 KO cells show no significant changes in fuel oxidation compared to the WT cells (Figure 8D).

**Figure 8.**
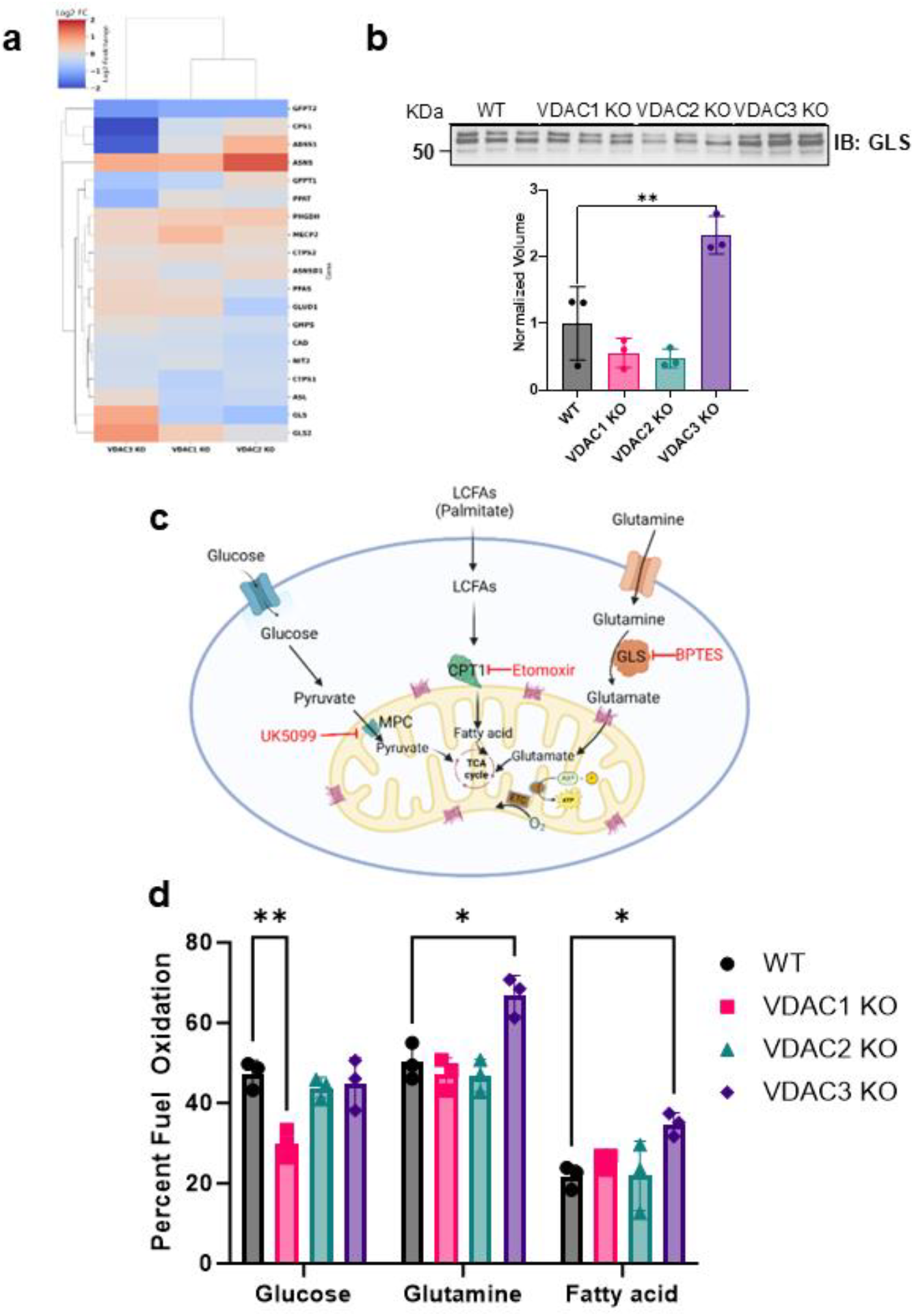
VDAC3 KO leads to increased reliance on glutamine metabolism. a) Heatmap of altered proteins associated with glutamine metabolism in VDAC1, VDAC2 and VDAC3 KO cells. b) Western blot of GLS expression in WT HeLa cells and VDAC isoform KO cell lines and the corresponding densitometry quantification. The bar graph represents the average of 3 repeats. c) Simplified metabolic map of mitochondrial fuel pathways and inhibitors used in Agilent Seahorse XF mitochondria fuel flex test (created on BioRender). d) Bar graph comparing the glucose, glutamine, and fatty acid dependency for WT (gray), VDAC1 KO (pink), VDAC2 KO (teal), and VDAC3 KO (purple). Data from 3 independent experiments are represented. The symbols represent data from independent experiments, and error bars indicate the standard deviation from the mean. Significance was tested using one-way ANOVA followed by the Dunnett post hoc test (*p < 0.05, **p < 0.01).

## Discussion

We characterized stable KOs of the three VDAC isoforms in HeLa cells. Knocking out both alleles of each VDAC isoform in an identical cellular background enabled us to compare isoform-specific effects across a battery of different assays.

VDAC1 KO decreased basal glycolysis and decreased mitochondria-bound HK1 and HK2. This is consistent with mice KO studies where VDAC1 KO impaired glucose tolerance and reduced mitochondria-bound HK activity ^55^. VDAC1 KO in H9C2 cells also decreased HKII bound to mitochondria even though it did not affect basal glycolysis ^47^. The different effects on glycolysis could be due to increased glycolysis in HeLa cells compared to H9C2 cells, which prefer oxidative phosphorylation similar to primary cardiomyocytes ^56^. HeLa cells also express a 10-fold increase in HKII compared to HEK-293 cells ^57^, which has been implicated in the increased glycolysis (Warburg effect) characteristic of cancer cells. In our experiment, VDAC1 KO did not affect mitochondrial respiration in HeLa cells in contrast to VDAC1 KO in HAP1 cells, which decreased mitochondrial respiration and reserve capacity by increasing complex-I-linked respiration ^46^. This discrepancy may be due to distinct responses to VDAC1 KO in different cell types. Comparing the effect of VDAC1 KO in two oxidative muscles (ventricle and soleus) and a glycolytic muscle (gastrocnemius) showed opposite effects on MOM permeability, which is dependent on VDAC, measured as apparent K_m_ [ADP] in different muscle types ^58^. The ventricle and gastrocnemius muscles showed an increase in K_m_ [ADP] compared to the soleus, which had decreased K_m_ [ADP]. Only the soleus muscle showed a decreased rate of respiration in the presence of maximum ADP (V_max_). This suggests that while VDAC1 KO only affects the MOM permeability in the ventricle and gastrocnemius muscles, both mitochondrial membranes are affected by VDAC1 KO in the soleus. More studies are needed to understand the basis for these differences.

Interestingly, VDAC2 KO does not strongly affect metabolism in our assays despite decreasing cell growth and altered mitochondrial proteome. This may be due to a compensatory upregulation of certain ETC complexes in proteomics studies. The decreased cell growth in VDAC2 KO may be due to changes in apoptosis. Proteomics data show changes in apoptosis-related proteins, such as down-regulation of Bak and upregulation of PYCARD in VDAC2 KO cells. This is consistent with past studies highlighting VDAC2’s role in apoptosis through complexation with Bak and BAX ^21,22,31,51,59,60^. Further studies are needed to confirm that the decreased cell growth in VDAC2 KO cells is related to changes in apoptosis pathways in HeLa cells.

Finally, to our surprise, VDAC3 KO decreased cell growth and showed a significant decrease in metabolic activity measured using MTS assay, confirming the decrease in reducing agents such as NADH and FAD measured by flow cytometry. VDAC3 KO decreases mitochondrial ATP and shows severe defects in mitochondrial respiration. Proteomic studies further confirmed the large-scale downregulation of mitochondrial proteins, such as ETC complexes involved in respiration for VDAC3 KO. We also found the upregulation of glutamine metabolism pathway enzymes in accord with increased glutamine dependence in VDAC3 KO cells.

Our results suggest an evolutionarily conserved metabolite transport function for the oldest VDAC3 isoform, given the decrease in mitochondrial respiration, increased glutamine metabolism, and the complete loss of spare respiratory capacity in VDAC3 KO cells. Spare respiratory capacity corresponds to cells’ ability to increase mitochondrial respiration during high energy demand. VDAC3 KO in mice results in mitochondrial dysfunction in sperm and heart, two of the most energy-demanding cells. VDAC3 KO-induced male infertility could be expected, given the high expression of VDAC3 in the testis ^24^. However, VDAC3 is only a minor isoform in the heart, and the expression profile of VDAC isoforms is similar to that of HeLa cells^61^. Notably, VDAC3 KO mouse heart muscles showed an increased apparent affinity for mitochondria to ADP (K_m_ [ADP]). In contrast, gastrocnemius, a mixed glycolytic/oxidative muscle, did not show significant changes in K_m_ [ADP] ^40^. This suggests cell type-specific changes in MOM permeability in VDAC3 KO mice.

One of the possible explanations for the different roles of VDAC isoforms in cell metabolism could be their different regulation by cytosolic protein partners such as hexokinase, tubulin, α-synuclein, and Bcl2 family proteins. Seminal work by Valdur Saks’ group showed that isolated mitochondria have lower K_m_ [ADP] than permeabilized cells ^62^. This effect was initially linked to cytoskeletal proteins and subsequently shown to be due to free tubulin blocking VDAC^3^. The addition of free tubulin to isolated mitochondria increased the K_m_ [ADP] of mitochondria, suggesting decreased permeability of MOM to ADP due to VDAC reversible blockage by dimeric tubulin. A second component with unchanged K_m_ [ADP] was identified, possibly due to a fraction of VDAC remaining open to ADP. This led to the hypothesis that two rates of ADP uptake could be due to differences in the binding affinity to free tubulin of the three VDAC isoforms. Later, Queralt-Martin et al. confirmed in experiments with VDAC reconstituted into planar membranes that tubulin (and α-synuclein) blocks VDAC1 with ∼100 times higher affinity than VDAC3 ^20^. This suggests that VDAC3 may always be open for metabolite transport in and out of mitochondria, while VDAC1 and VDAC2 are mostly blocked by one of their cytosolic partners, such as tubulin or α-synuclein. This is consistent with calculations by Marko Vendelin’s group suggesting that only 2% of VDAC channels are open (accessible for cytosolic ADP) in cardiomyocytes ^63^. Finally, VDAC3 knock-down in HepG2 cancer cells also resulted in the largest decreased ATP, ADP, and NAD(P)H compared with knock-down of the other two VDAC isoforms^64^. These results obtained in HepG2 cells were also interpreted as VDAC1 and VDAC2 being mostly closed by free tubulin and VDAC3 being less sensitive to tubulin. Interestingly, theoretical calculations predicted that VDAC3 may prevent electrical suppression of MOM permeability at elevated MOM potential^65^. Overall, these results allow us to speculate that the least expressed VDAC3 isoform is constitutively open in cancer cells for essential metabolite transport function.

Alternatively, VDAC3 expression may constitute part of a signaling axis that promotes oxidative phosphorylation similar to VDAC2’s functions in apoptosis, not through direct channel properties but through the recruitment of Bak. VDAC3 may be a critical signaling intermediate via protein-protein interactions that ultimately promote nuclear expression of pro-OXPHOS gene expression through PGC1-μ or an alternative pathway ^66^. To test this hypothesis, future studies are needed to determine the interacting partners unique to VDAC3 using immunoprecipitation-mass spectrometry. Additionally, following the approach performed for VDAC2’s binding to Bak^60^, VDAC2/VDAC3 chimeras can be synthesized to determine which regions of the channel are critical for maintaining the respiration phenotype.

In conclusion, comparing the role of each VDAC isoform in a single cell type allowed us to reveal the distinct roles of each VDAC isoform to be revealed. Consistent with previous studies, we show that VDAC1 and VDAC2 are involved in glycolysis and apoptosis, respectively. To our surprise, VDAC3, the oldest but minor VDAC isoform, which was initially assumed not to form a channel, plays the most crucial role in regulating mitochondrial function and metabolic pathways in HeLa cells. We propose that VDAC3 uniquely enables cells to sustain higher energy demands, explaining its role in energy-demanding cell types such as the heart and sperm. Determining whether VDAC3’s effect on metabolism is direct or happens via a yet undescribed cellular signaling pathway represents the next avenue for interrogation. Future studies are needed to understand clonal variability in CRISPR-Cas9 KO cell lines and the role of the VDAC3 isoform in cancer metabolism.

## Data Availability

The raw and processed proteomics data are available in the MassIVE database under the identifier MSV000097150. Any additional information required to reanalyze the data reported in this paper is available from the authors upon request. All other data are available in the main text or Supplementary information.

## Acknowledgments

M.R., W.F., B.G.B., N.A.B., T.K.R. and S.M.B. were supported by the Intramural Research Program of the Eunice Kennedy Shriver National Institute of Child Health and Human Development of the National Institutes of Health. W.M.R. was supported by the National Institute of Neurological Disorders and Stroke, NIH, and a generous grant from the J. Yang and Family Foundation, Caltech.

## Author Contributions

M.R., W.M.R., T.K.R., and S.M.B. designed the experiments; M.R. performed and analyzed Seahorse real-time cell metabolic assays; W.M.R. and B.Q. performed and analyzed mass spectrometry experiments; W.F. performed and analyzed flow cytometry assays; B.G.B. and M.R. performed and analyzed microscopy images; M.R., W.M.R., D.H., J.H., and N.A.B. performed and analyzed western blot experiments; M.R., W.M.R., T.K.R., and S.M.B. wrote the manuscript; all authors contributed intellectually, revised, and edited manuscript.

## Supplementary Information

Supplementary information accompanies the manuscript.

