## Supplemental Material for "Unlocking the Distinct Roles of the Three Mammalian VDAC Isoforms in Mitochondrial Respiration and Cancer Cell Metabolism"

### Supplementary Methods

##### Flow cytometry

Cells were seeded at 2x$10^5$ cells/well in 12 well plates and incubated overnight at 37 °C and 5% $CO_2$. The cells were harvested by trypsinization, washed with PBS, and centrifuged. Cell pellets were resuspended and stained with Live/Dead Near-IR stain (ThermoFisher Scientific, L34976) for 15 mins, washed with PBS, and centrifuged. The cell pellets were resuspended in 250 µL of PBS and immediately acquired on a BD FACSymphony A5 flow cytometer (BD Biosciences). The fluorescence of NADH and FAD was measured by exciting the cells with a 355 nm UV laser, and the emission was detected with bandpass filters of 580/20 nm (FAD) and 450/50 nm (NADH). Data were analyzed with FlowJo v10.8 (BD Biosciences) with gating on live single cells, and NADH and FAD were measured as mean fluorescence intensity.

##### ATP Production Rate Assay

Cells were seeded at 10-15x$10^3$ cells/well in XFe96/XF Pro PDL cell culture microplates (Agilent, 103798) and incubated overnight at 37 °C and 5% $CO_2$. The following day, media was replaced with Seahorse XF DMEM Media, pH 7.4 (Agilent, 103575) supplemented with 10 mM glucose (Agilent, 103577), 1 mM pyruvate (Agilent, 103578), and 2 mM glutamine (Agilent, 103579). ATP rate assay (Agilent, 103591) was performed using an XF Pro analyzer (Agilent) according to the manufacturer's protocol. Hoechst 33342 (ThermoFisher Scientific, 62249) was added to the last port to stain nuclei for cell count and imaged using Cytation5 (BioTek).

##### Western blot for VDAC isoforms.

Cells were seeded at 1x$10^6$ cells/well in 6 well plates and incubated overnight at 37 °C and 5% $CO_2$. The cells were harvested by trypsinization, washed with PBS, and centrifuged. Cell pellets were resuspended in 100 µL lysis buffer (50 mM Tris-HCl pH 7.5, 150 mM NaCl, 5mM EDTA, 1% Triton X-100) with cOmplete, EDTA-free protease inhibitor cocktail (Sigma Aldrich, 4693132001), and incubated on ice for 10 min with occasional vortex. Samples were centrifuged at 14,000 rpm at 4 °C for 10 min, and the supernatant was transferred into a new 1.5 mL tube. Total soluble protein concentrations were measured using the Qubit™ Protein BR Assay (ThermoFisher Scientific, A50668). Samples were mixed with NuPAGE™ LDS Sample Buffer (4X) (ThermoFisher Scientific, NP0007) and 5% 2-mercaptoethanol (Fisher BioReagents, BP176) and heated at 95 °C for 10 min. Equal amounts of protein samples were loaded and separated using NuPAGE™ Bis-Tris Mini Protein Gels, 4–12% (ThermoFisher Scientific, NP0323BOX) and transferred to nitrocellulose membranes using iBlot™ 2 Transfer Stacks (ThermoFisher Scientific, IB23001). Membranes were blocked with 3% w/v BSA prepared in TBST buffer for 30 min at room temperature, incubated with primary antibodies overnight at 4 °C, washed with TBST (three times for 10 min each), incubated with appropriate secondary antibodies for one hour at room temperature, and washed with TBST buffer (six times for 5 min each). The blots were incubated in Pierce™ ECL Plus Western Blotting Substrate (ThermoFisher Scientific, 32134) for 5 min and imaged using the iBright FL1500 imaging system (ThermoFisher Scientific).

| **ANTIBODIES** | **SOURCE** | **IDENTIFIER** |
|---|---|---|
| Anti-VDAC1/Porin + VDAC3 antibody [20B12AF2] | Abcam | ab14734 |
| Anti-VDAC2 antibody | Abcam | ab37985 |
| Anti-VDAC3 antibody | Abcam | ab130561 |
| Donkey Anti-Mouse IgG H&L (HRP) preadsorbed | Abcam | ab7061 |
| Donkey Anti-Goat IgG H&L (HRP) | Abcam | ab97110 |
| Donkey Anti-Rabbit IgG H&L (HRP) | Abcam | ab205722 |

### Supplementary Figures

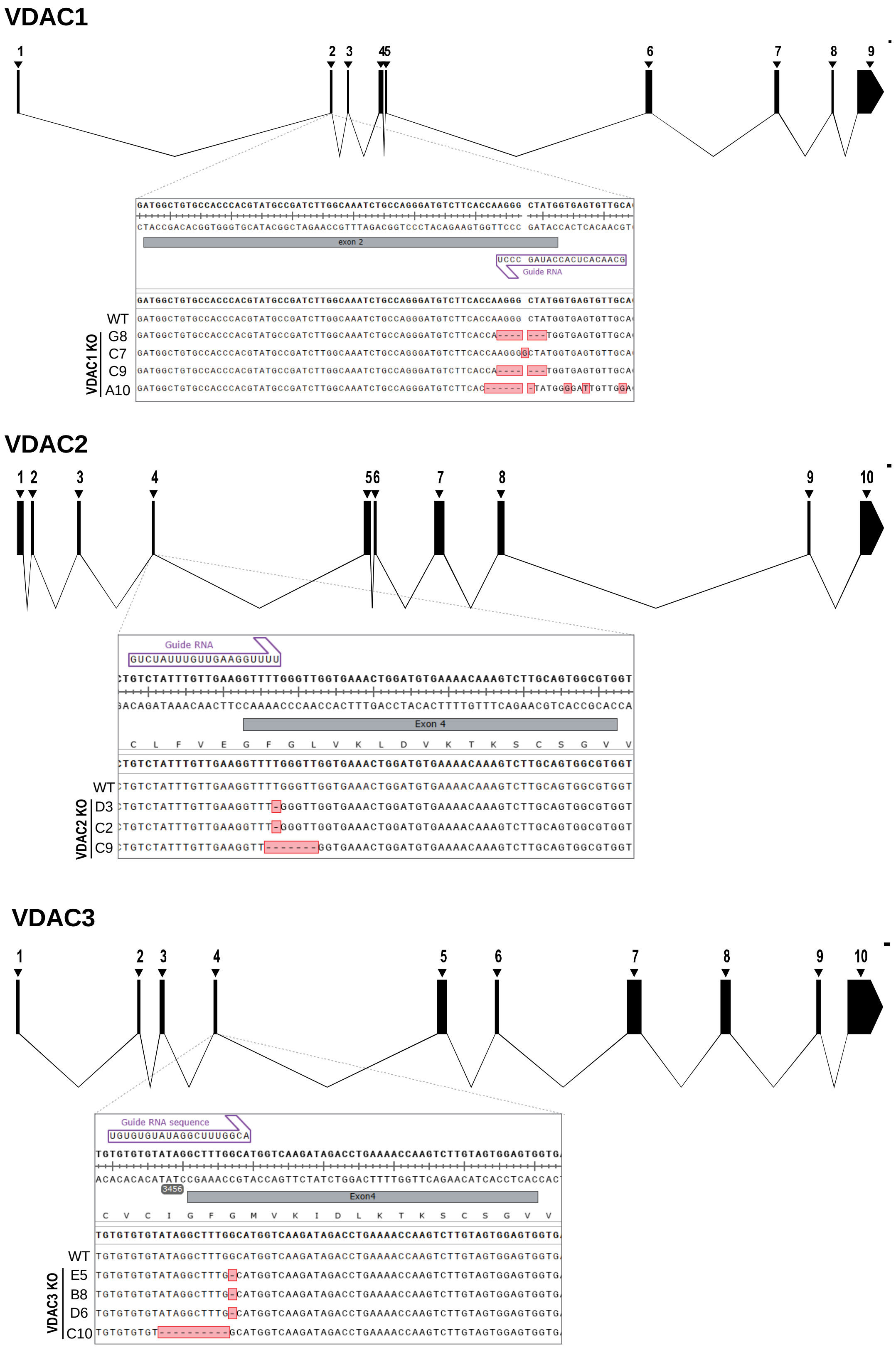

**Supplementary Figure S1.** Human genomic maps of VDAC isoforms, the sites of guide sgRNA, and the alignment of DNA sequencing results confirming VDAC1, 2, and 3 KO indels in HeLa cells.

a

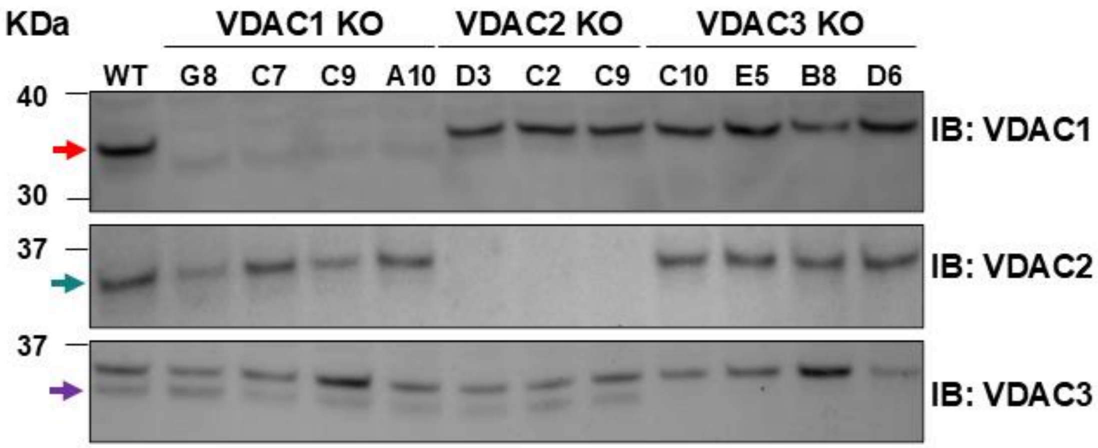

b

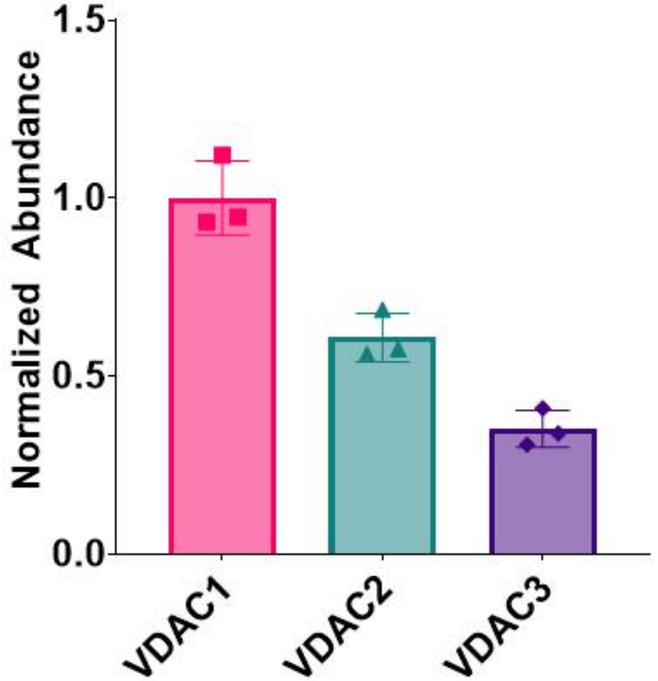

**Supplementary Figure S2. VDAC1, 2, and 3 expression in HeLa cells** A) Western blot VDAC1, VDAC2, and VDAC3 expression levels in HeLa WT and KO clones. The top band (pink arrow) corresponds to VDAC1 expression in VDAC1 immunoblot (IB), and the bottom band (purple arrow) corresponds to VDAC3 expression in VDAC3 IB. B) VDAC isoform expression ratio measured by mass spectrometry in WT HeLa cells normalized to VDAC1 expression level.

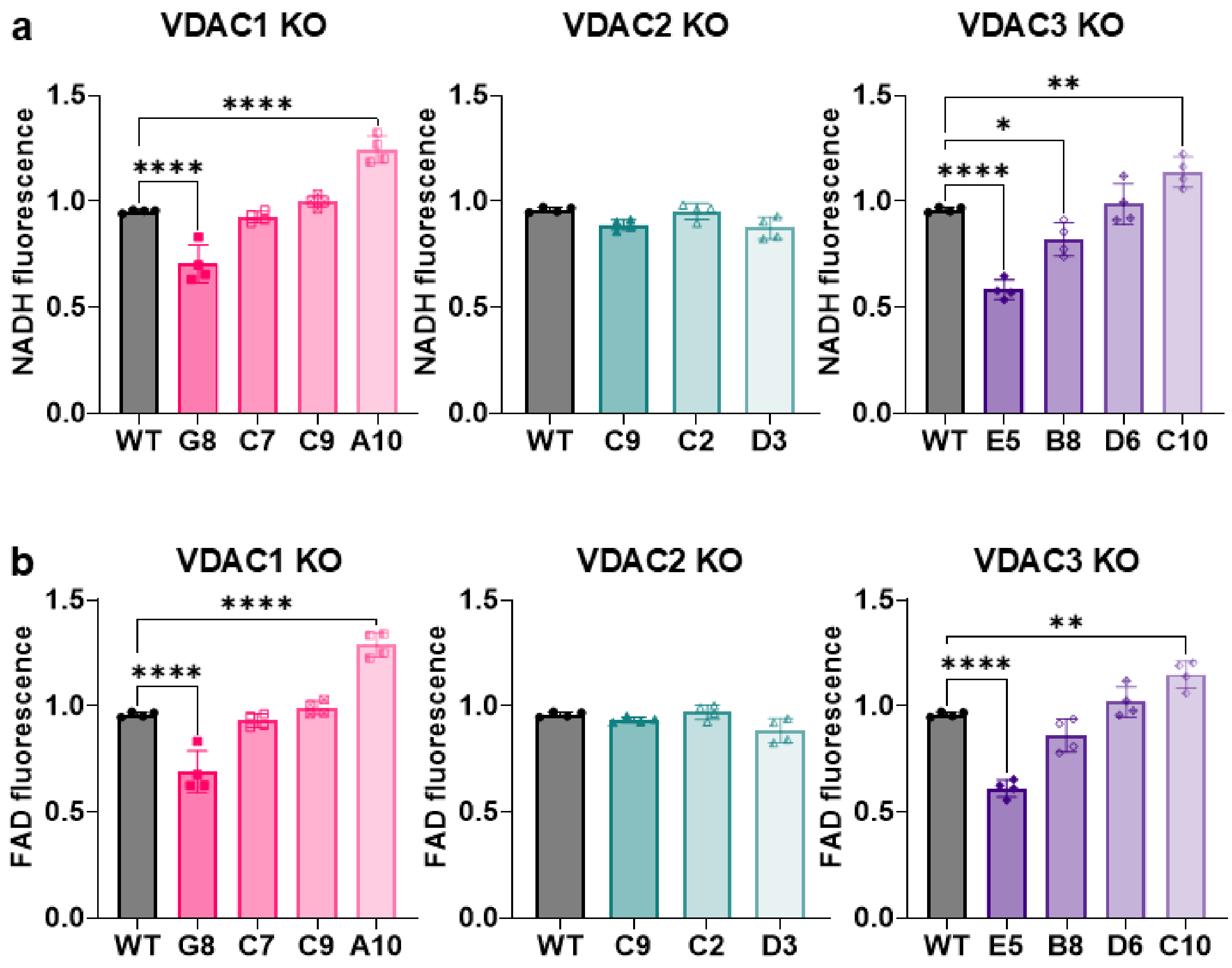

**Supplementary Figure S3. NADH and FAD autofluorescence in HeLa WT and VDAC isoform KO cells.** Normalized NADH fluorescence (A) and FAD fluorescence (B) in VDAC1, VDAC2, and VDAC3 KO clones compared to HeLa WT (gray) were measured using flow cytometry (see Supplemental Methods). While VDAC2 KO clones do not affect NADH and FAD fluorescence, VDAC1 and VDAC3 KOs show clonal variability in NADH and FAD fluorescence. Clones VDAC1 KO G8 and VDAC3 KO E5 show the most significant decrease in NADH and FAD. The symbols represent data from four independent experiments. Error bars indicate the standard deviation from the mean. Significance was tested using one-way ANOVA followed by the Dunnett post hoc test (*p<0.05, **p<0.01, ****p<0.0001).

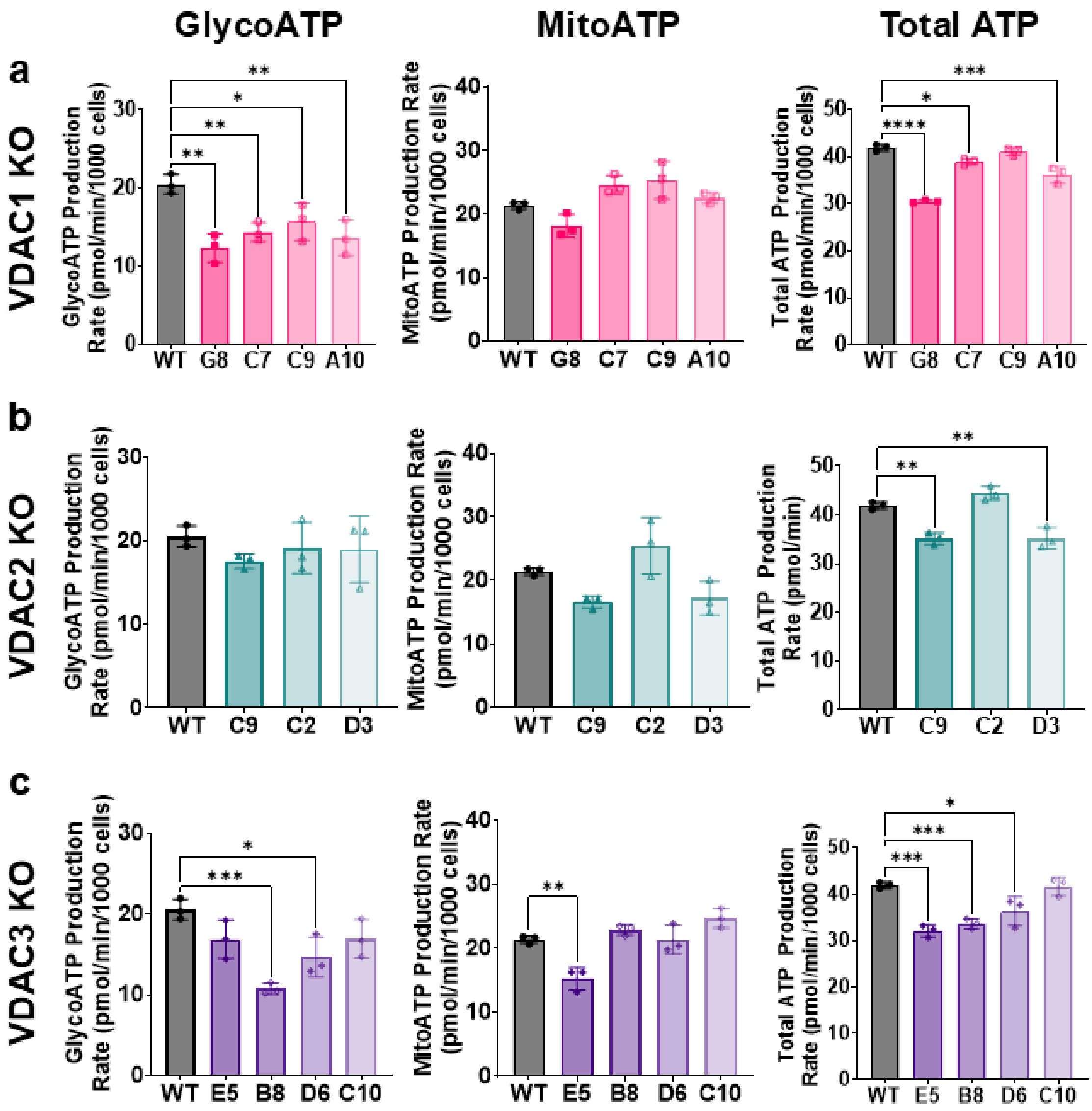

**Supplementary Figure S4. ATP production rate in HeLa WT and VDAC isoform KO cells**. The total, glycolytic (GlycoATP), and mitochondrial (MitoATP) ATP production rate in HeLa cells with VDAC1 KO (A), VDAC2 KO (B), and VDAC3 KO (C) clones. A) All VDAC1 KO clones show a significant decrease in glycoATP production rate compared to the WT. There is no significant difference in mitoATP production rate, but the minor differences result in a variable total ATP production rate. VDAC1 KO G8 clone has the largest decrease in total ATP production rate. B) GlycoATP and mitoATP are not significantly affected in VDAC2 KO clones, while the total ATP production rate is decreased by ~16% for VDAC2 KO C9 and D3 clones. C) VDAC3 KO results in variable glycoATP and mitoATP production rates for each clone. GlycoATP production rate is significantly decreased for VDAC3 KO clones B8 (~48%) and D6 (~29%). Only VDAC3KO E5 shows a significant decrease in mitoATP production rate. This results in some variability of the total ATP production rate, with the VDAC3 KO E5 clone having the largest effect on the total ATP production rate. The symbols represent data from three independent experiments. Error bars indicate the standard deviation from the mean. Significance was tested using one-way ANOVA followed by the Dunnett post hoc test (* $p < 0.05$, ** $p < 0.01$, *** $p<0.001$, **** $p<0.0001$).

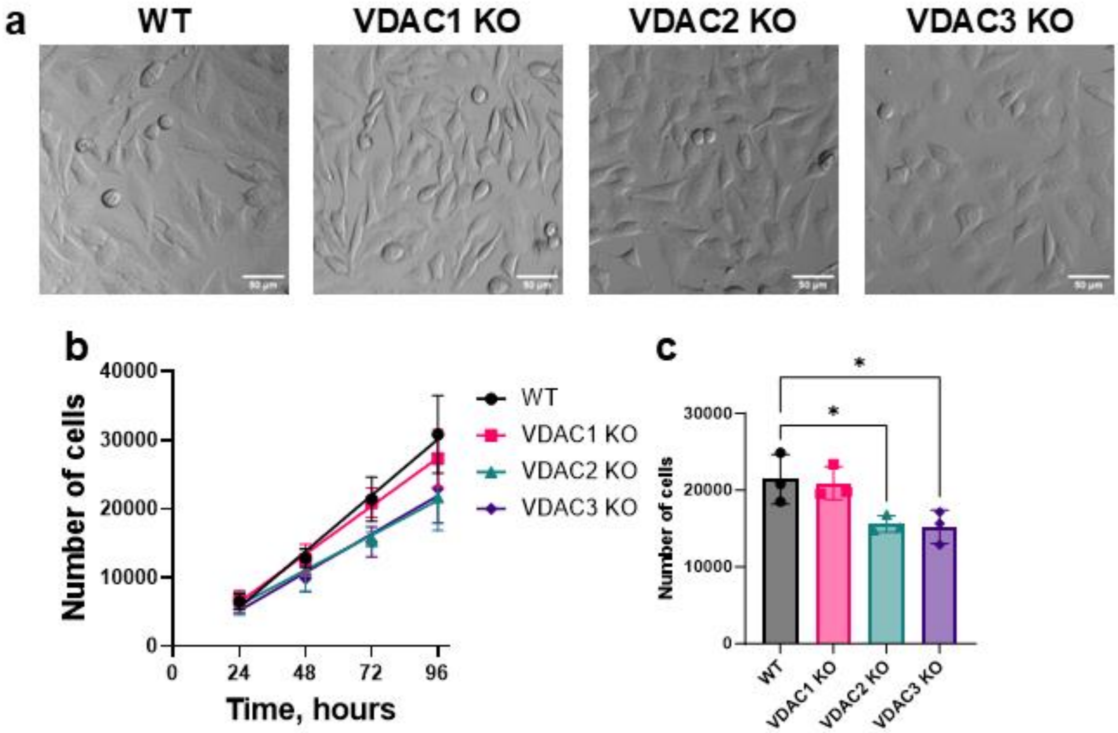

**Supplementary Figure S5. Cell morphology and growth of HeLa WT and VDAC isoform KO cells** A) Brightfield images of HeLa cells WT, VDAC1 KO, VDAC2 KO, and VDAC3 KO. B) Number of cells measured using Hoechst 33342 staining every 24 hours and C) bar graph represents the number of cells after 72 hours for WT (gray circle), VDAC1 KO (pink square), VDAC2 KO (teal triangle), and VDAC3 KO cells (purple diamond). The symbols represent data from three independent experiments, and error bars indicate the standard deviation from the mean. Significance was tested using one-way ANOVA followed by the Dunnett post hoc test (*p< 0.05).

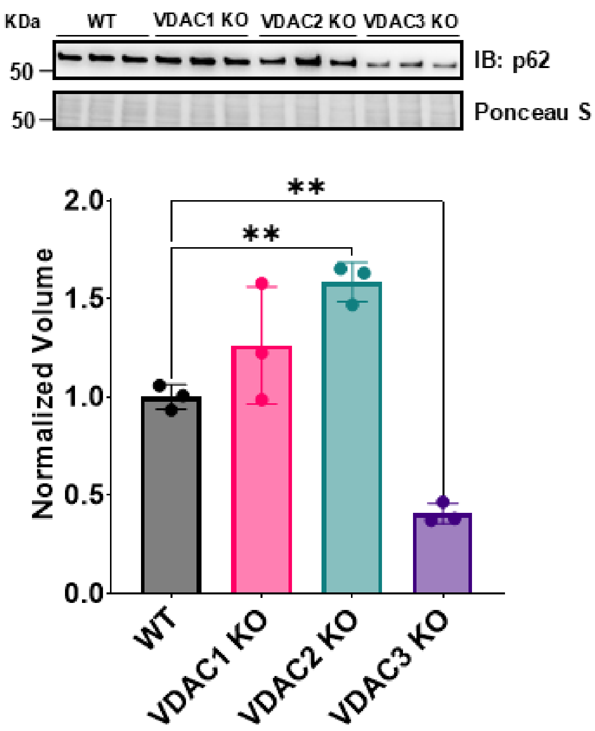

**Supplementary Figure S6. p62 expression in HeLa WT and VDAC isoform KO cells.** Western blot and densitometry quantification of p62 expression in WT HeLa and VDAC isoform KO cell lines. The symbols represent data from three independent experiments, and error bars indicate the standard deviation from the mean. Significance was tested using one-way ANOVA followed by the Dunnett post hoc test (**$p < 0.01$).

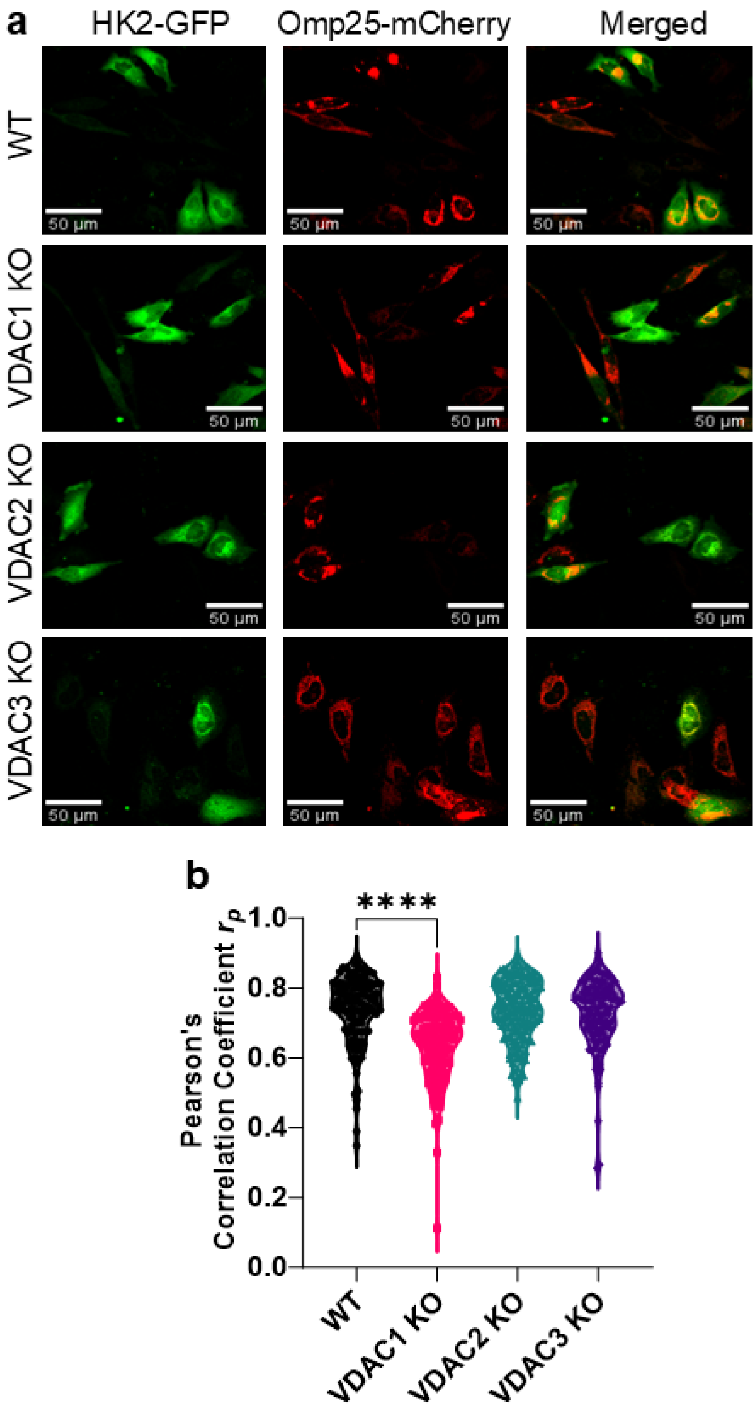

**Supplementary Figure S7. Mitochondrial localization of HK2 in HeLa WT and VDAC isoform KO cells.** A) Representative images of HK2-GFP (green) colocalization with Omp25-mCherry (red) shown in the merged image (yellow) for HeLa cells WT, VDAC1 KO, VDAC2 KO, and VDAC3 KO. B) Colocalization analysis of HK2 with Omp25 measured by Pearson's correlation coefficient (PCC) shows a significant decrease in HK2 mitochondrial localization in VDAC1 KO. Data from three independent experiments are represented. The symbols represent PCC for each cell. Significance was tested using one-way ANOVA followed by the Dunnett post hoc test (****$p<0.0001$).
